# An engineered Clostridial mini consortium modulates intestinal inflammation

**DOI:** 10.64898/2026.08.11.743107

**Authors:** Edward Ionescu, Jack H. Arnold, Christopher R. Weber, Mark Mimee, Cathryn R. Nagler

## Abstract

Modern lifestyle factors have altered gut microbiota composition and function. Bacteria in the Clostridia class modulate mucosal immune responses through various mechanisms including production of secondary bile acids (SBA). Here, we present a novel system to study how the SBA isodeoxycholic acid (isoDCA) regulates host immunity. Through targeted mutagenesis of bile acid epimerization genes, we engineered *Ruminococcus gnavus* to ablate isoDCA production. Combining *R. gnavus* (WT or KO) with *Peptacetobacter hiranonis* created a two-member consortium that toggles isoDCA production on or off while keeping all other variables constant. Using this system, we demonstrate that isoDCA induces colonic lamina propria RORγt□ Foxp3□ regulatory T cells (pTregs) through a mechanism requiring both the Takeda G protein-coupled receptor 5 (TGR5) and the Farnesoid X receptor (FXR). Engraftment of this isoDCA+ consortium protected against colitis in an adoptive T cell transfer model by reshaping the microbiota and suppressing host inflammation.

## Introduction

21^st^ century lifestyle practices have altered the composition and function of the commensal microbiota. We, and others, have linked these changes to the increasing prevalence of immune-mediated, noncommunicable chronic diseases (NCCDs) including food allergies, inflammatory bowel diseases, obesity, asthma and diabetes^1–3^. Resident gut microbes are intimately intertwined with the gut associated lymphoid tissues, which contain one of the largest populations of immune cells in the body^4,5^. Despite constant interaction with trillions of potentially inflammatory microbes, a healthy intestinal immune system must remain non-responsive toward harmless antigens from commensal bacteria and food while maintaining the capacity for rapid, robust responses to pathogens. Lachnospiraceae, an abundant bacterial family in the human (and mouse) gut, are generally considered to contain health promoting bacteria^6,7^. In earlier work we have shown that mucosa-associated bacteria in the Clostridia class (which contains this family) regulates sensitization to food allergens^8^. To gain insight into the mechanisms responsible for this bacteria-induced barrier protective response, we have examined the roles of various Lachnospiraceae products including the short chain fatty acid butyrate^9–11^, tryptophan metabolites and commensal flagellins^12^. Here we extend this analysis to an examination of the immunomodulatory capacity of Lachnospiraceae-derived secondary bile acids (SBAs). Bile acids (BAs) are commonly known as emulsifying agents which facilitate the absorption of dietary lipids in the small intestine^13^. BAs also act in a hormone-like fashion to regulate their own biosynthesis as well as glucose homeostasis^14^. Primary bile acids (PBAs), produced in the liver from the metabolism of cholesterol, are stored as taurine or glycine conjugated bile salts in the gall bladder^13^. PBAs are released into the duodenum after a meal; 95-97% of PBA are reabsorbed in the ileum. The remaining 3-5% continue to the colon where intestinal bacteria convert them into SBAs; almost all of the BA pool in the colon has therefore undergone modification by commensal bacteria. The first modification of BAs, deconjugation of taurine or glycine, is performed by bile salt hydrolases (BSH), which are conserved across the major gut bacterial phyla^13^. The conversion of PBAs to SBAs is a multistep pathway mediated by the 7α-dehydroxylation enzymes encoded by the *bile acid inducible* (*bai*) operon^15^. Most of the small number of *bai*-expressing taxa identified to date are in the Clostridia class^16–18^. Although *bai* expressing bacteria are rare, and at low abundance, they process high concentrations (about 1mM^19,20^) of colonic PBA and, in so doing, exert a considerable impact on the overall bacterial metabolite pool. The most abundant SBA end products of this pathway, deoxycholic acid (DCA) and lithocholic acid (LCA), can undergo further modifications like epimerization by 3a-and 3b-hydroxysteroid dehydrogenases (HSDH). DCA, LCA and their derivatives comprise over 90% of the fecal BA pool^15^. Potent immunoregulatory properties have been described for two modified SBA, isoDCA and isoalloLCA ^21–23^. The detergent activity of many SBAs can also disrupt membrane integrity to selectively lyse or inhibit the growth of intestinal bacteria and shape the composition and function of the intestinal microbiota^13,24^.

The recalcitrance of bacteria in the Clostridia class to genetic manipulation has made it difficult to demonstrate a direct mechanistic role for Clostridial metabolites in promoting intestinal homeostasis. This resistance is driven by a lack of compatible replication origins, inefficient homologous recombination, and robust endogenous defense systems (like restriction modification systems) which actively degrade exogenous DNA^25–28^. In recent years, however, advancement of genetic tools for engineering non-model Clostridia have been described^19,25,29–31^. In this manuscript we describe a novel, engineered strain of *Mediterraneibacter [Ruminococcus] gnavus* with ablated production of the immunomodulatory SBA isoDCA. By combining *Peptacetobacter hiranonis* with this engineered strain of *R. gnavus,* or the wild-type (WT) strain, we created two isogenic consortia of Clostridia that can turn isoDCA production “on” or “off”. This system uniquely enabled investigation into how isoDCA, produced by native species from physiologically relevant concentrations of host-derived taurocholic acid (TCA), promotes intestinal homeostasis. Engraftment of antibiotic-treated SPF mice with each of the consortia revealed that isoDCA can induce both ileal and colonic lamina propria (LP) RORγt^+^Foxp3^+^ pTregs. Leveraging this system’s ability to isolate the function of a single microbial metabolite in mice with a replete microbiome we identified a previously unknown requirement for the Takeda G protein-coupled receptor 5 (TGR5), and to a lesser extent, FXR, in isoDCA-mediated induction of colonic LP RORγt^+^Foxp3^+^ pTreg. Lastly, we demonstrated that our isoDCA-producing (isoDCA+) consortium protects from colitis in the CD45RB^hi^ adoptive T cell transfer model. No protection was mediated by the isogenic, isoDCA-deficient (isoDCA-) control consortium. Taken together, this work defines a novel approach for studying a prominent gut metabolite, highlighting its immunosuppressive effects and importance for the maintenance of intestinal homeostasis.

## Results

### Engineering isogenic consortia to interrogate the immunomodulatory effects of isoDCA

To better study the immunomodulatory effects of SBAs, we first engineered an isogenic strain of Clostridia to ablate isoDCA production. *R. gnavus* is an abundant species of Clostridia that converts the predominant SBA deoxycholic acid (DCA) into isoDCA^24^. IsoDCA is the 3β-epimer of DCA and is generated through a two-step epimerization mediated by 3α-and 3β-HSDHs (**Figure 1A**). To ablate isoDCA production in *R. gnavus*, we designed a plasmid (pJHA342) to functionally knock-out the 3β-HSDH required for converting 3oxo-DCA into isoDCA, the terminal enzymatic step (**Figure 1B**). This plasmid encoded a group II intron, designed by the ClosTron intron design tool ^29^, to disrupt the coding sequence of *rumgna_00694* (3β-HSDH gene) in *R. gnavus*. The intron first initiates a double strand DNA break in *rumgna_00694* to target a 5’ nucleotide insertion of RNA and then reverse transcribes the single strand RNA into DNA via the LtrA recombinase (reverse transcriptase) (**Figure 1B**). After host DNA repair, the group II intron ultimately disrupts *rumgna_00694* gene function due to the nucleotide insertion. This ∼1kb DNA insertion was confirmed via PCR using primers flanking *rumgna_00694* (**Figure 1C**). With this strategy, we engineered a novel strain, *R. gnavus* Ω*rumgna_00694* (*R. gnavus* KO). To further characterize this engineered strain, *R. gnavus* KO, we first sought to confirm whether our gene disruption resulted in a functional knock-out of the 3β-HSDH-mediated reduction of 3oxo-DCA required for isoDCA production. Culturing *R. gnavus* WT and *R. gnavus* KO *in vitro* with DCA and analyzing bile acids in culture supernatants via thin layer chromatography (TLC) confirmed that *R. gnavus* WT could produce isoDCA from DCA while *R. gnavus* KO lost isoDCA production (**Figure 1D**), highlighting that the group II intron mutagenesis successfully ablated isoDCA production. Minimal production of 3oxo-DCA was observed in cultures of *R. gnavus* KO with DCA (**Figure 1D**).

**Figure 1.**
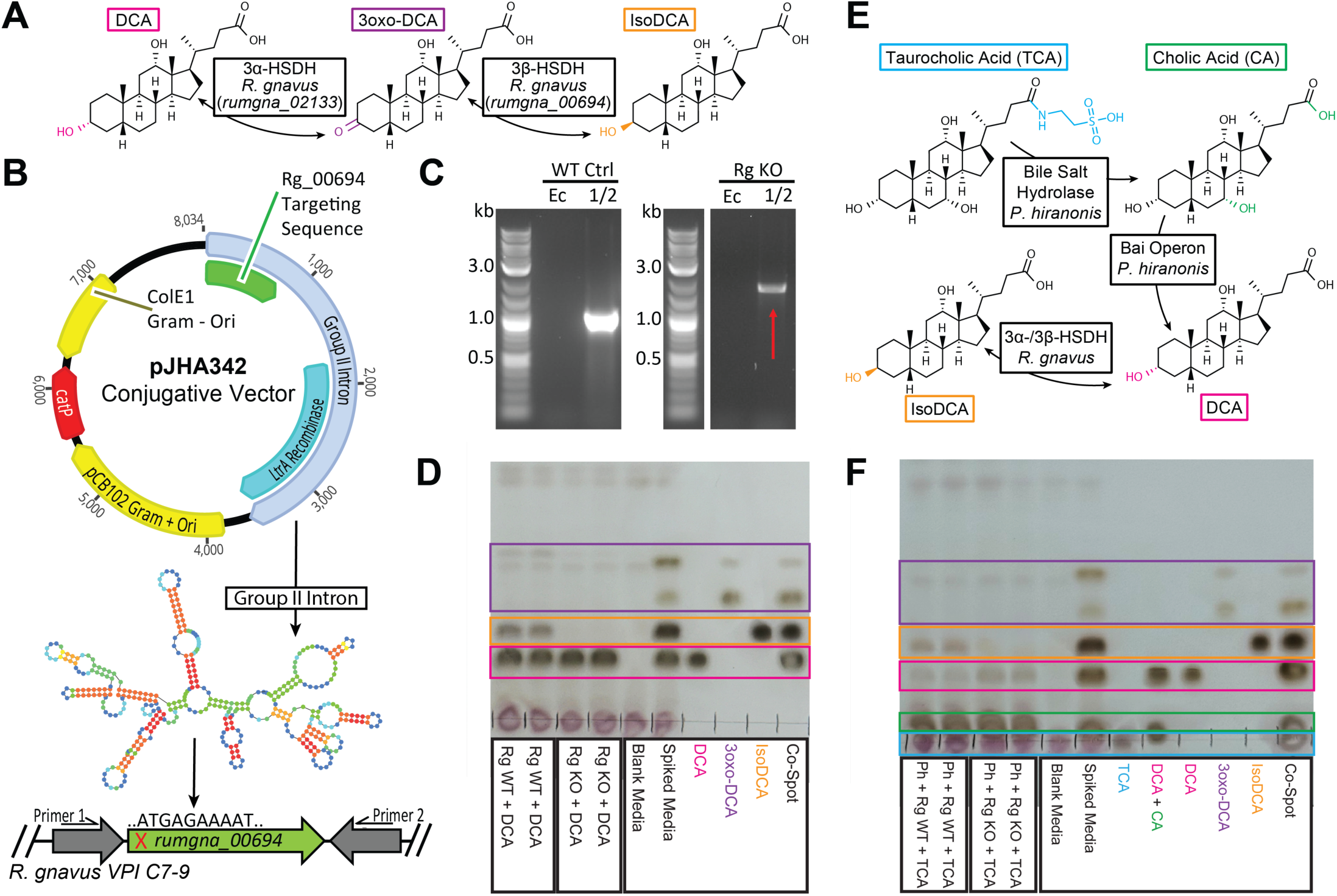
An engineered consortium of *Peptacteobacter hiranonis* and *Ruminococcus gnavus Δrumgna_00694* cannot produce isoDCA *in vitro*. (A) Schematic of the 2-step epimerization of DCA into isoDCA by *R. gnavus*. (B) Group II intron-based approach to engineer *R. gnavus*. (C) PCR of *rumgna_00694* in *R. gnavus* WT (Rg WT) and *R. gnavus* Ω*rumgna_00694* (Rg KO). Ec = *Escherichia coli* primers used for conjugation. 1/2 = primers for *rumgna_00694* as seen in (B). (D) Representative thin layer chromatography (TLC) of culture supernatants of Rg WT and Rg KO supplemented with DCA. (E) Schematic of the conversion of TCA into CA and DCA by *P. hiranonis* and into isoDCA by *R. gnavus*. (F) Representative TLC of culture supernatants of *P. hiranonis* (Ph) supplemented with TCA, then cultured with Rg WT or Rg KO. For (D, F), repeat lanes indicate biological replicates.

To confirm that the genetic manipulation did not alter the activity of the 3α-HSDH (*rumgna_02133*) required for the conversion of DCA to 3oxo-DCA, additional cultures of *R. gnavus* KO were grown with 3oxo-DCA and supernatants were analyzed by TLC. *R. gnavus* KO predominantly reverted the 3oxo-DCA substrate back into DCA, and did not generate isoDCA, which further confirmed ablation of isoDCA production in this strain (**Figure S1A**). For additional characterization of the genetic insertion, the novel *R. gnavus* KO strain was complemented using plasmid pJHA826 to inducibly express *rumgna_00694* in *R. gnavus* KO (**Figure S1B**)^32^. By culturing *R. gnavus* KO/pJHA826 with DCA and inducing *rumgna_00694* expression, isoDCA production was observed (**Figure S1C**), confirming that the ablation of isoDCA production in *R. gnavus* KO was indeed due to the disruption of *rumgna_00694* and not an off-target gene.

DCA is a microbially-derived SBA and thus, *R. gnavus* WT and *R. gnavus* KO alone are insufficient to study the immunomodulatory effects of isoDCA *in vivo*. To create an isogenic consortium for use *in vivo*, we paired *R. gnavus* with the Clostridial species *P. hiranonis,* which is one of the very few Clostridia that possesses both a BSH, for deconjugation of PBAs, and the intact *bai* operon, required for the 7α-dehydroxylation of cholic acid (CA) into DCA^33–36^ (**Figure 1E**). To verify that the consortia metabolized bile acids as expected, *P. hiranonis* was cultured *in vitro* with TCA and then with either *R. gnavus* WT or *R. gnavus* KO, and the supernatants were analyzed via TLC. As expected, both culture conditions yielded CA and DCA indicating effective deconjugation and 7α-dehydroxylation by *P. hiranonis* (**Figure 1F**). Additionally, *R. gnavus* WT, but not *R. gnavus* KO, could generate isoDCA under these conditions (**Figure 1F**). Taken together, these results indicate that a consortium of *P. hiranonis* and *R. gnavus* WT (isoDCA+ consortium) can produce isoDCA from a host-derived BA (TCA) and a consortium of *P. hiranonis* and *R. gnavus* KO (isoDCA-consortium) is an effective isogenic control to isolate the effects of isoDCA.

### IsoDCA regulates intestinal lamina propria Treg populations

To evaluate how isoDCA promotes intestinal homeostasis *in vivo* in a native context, the consortia needed to achieve sustained isoDCA production *in situ*. Therefore, it was necessary for the isoDCA+ and isoDCA-consortia to stably engraft in mice. Broad-spectrum antibiotics (ABX) are commonly used to transiently deplete the gut microbiota, reducing total bacterial burden and clearing competitive ecological niches in the cecum and colon^37–40,41^. After cessation of antibiotics, bacterial load recovers within a couple of weeks^40,42^. To facilitate the engraftment of the isoDCA+ and isoDCA-consortia, mice were treated neonatally and post-weaning with an ABX-cocktail consisting of ampicillin, vancomycin, neomycin, and metronidazole (AVNM) prior to administration of the consortia via seven days of intragastric (i.g.) gavage (**Figure 2A**). Abundance of both *P. hiranonis* and *R. gnavus* were checked before, during, and after the colonization period by qPCR with species-specific primers which confirmed engraftment in both groups at two weeks after colonization began (**Figures 2B and 2C**). Fecal bacterial load, measured via qPCR with universal 16S primers, returned to levels comparable to age-matched, untreated SPF mice by day 35 (**Figure 2D**).

**Figure 2.**
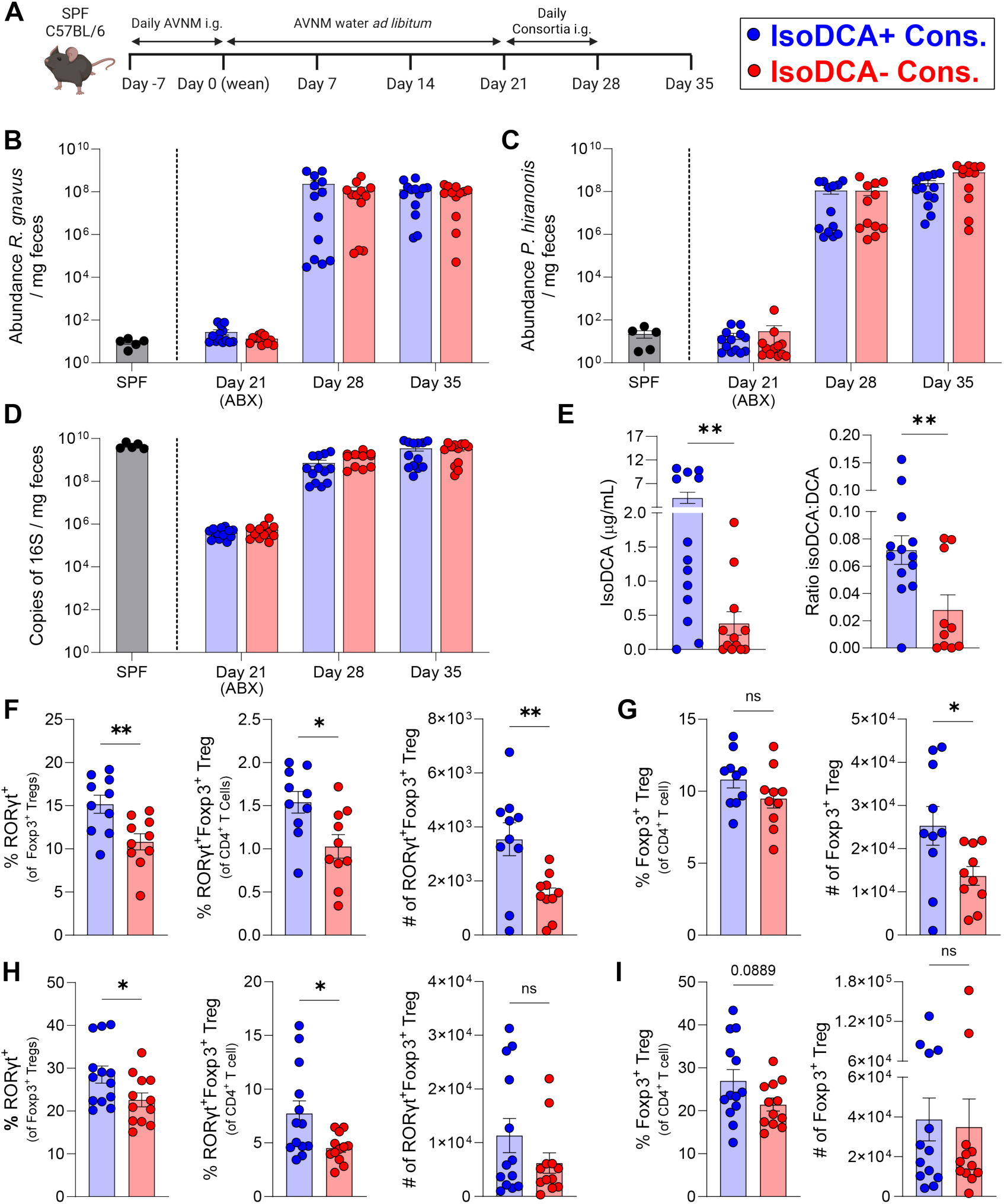
IsoDCA+ consortium colonizes antibiotic-treated mice and induces colonic lamina propria RORγt^+^FoxP3^+^ regulatory T cells. (A) Experimental timeline. (B-C) Abundance of *R. gnavus* (B) and *P. hiranonis* (C) normalized to fecal mass. (D) Fecal bacterial load normalized to fecal mass. (E) Fecal isoDCA concentration (left) and ratio of isoDCA to DCA fecal concentration (right) on day 35. (F) Proportion of RORγt^+^Foxp3^+^ Tregs of Foxp3^+^ Tregs (left) and of CD4^+^ T cells (center) and number of RORγt^+^Foxp3^+^ Tregs (right) in the ileal lamina propria (LP) on day 35. (G) Proportion (left) and number (right) of Foxp3^+^ Tregs in the ileal LP on day 35. (H) Proportion of RORγt^+^Foxp3^+^ Tregs of Foxp3^+^ Tregs (left) and of CD4^+^ T cells (center) and number of RORγt^+^Foxp3^+^ Tregs (right) in the colonic LP on day 35. (I) Proportion (left) and number (right) of Foxp3^+^ Tregs (right) in the colonic LP on day 35. For (B-D), SPF = age-matched SPF mouse (*n* = 5). Data in (B-I) are pooled from two independent experiments. For (B-D), *n* = 13 isoDCA+ consortium, 12 isoDCA-consortium. For (E), *n* = 13 isoDCA+ consortium, 12 isoDCA-consortium (10 isoDCA-consortium for ratio due to zero DCA concentration). For (F, G), *n* = 10 isoDCA+ consortium, 10 isoDCA-consortium. For (H, I), *n* = 13 isoDCA+ consortium, 12 isoDCA-consortium. For (B-I), symbols represent individual mice and error bars represent mean ± SEM. Statistics were analyzed by Student’s t-test. * *P* < 0.05, ** *P* < 0.01.

Since the isoDCA+ consortium and isoDCA-consortium are isogenic, they are expected to behave similarly with regards to their broad effects on microbiota composition. To examine this, 16S sequencing was performed on fecal samples of colonized mice collected on day 35 (**Figure S2A**). The microbiomes of the two groups were comparable in overall composition at the class level (**Figure S2B**). Analysis of different abundance taking sample and scale variation into account (ALDEx2, ref. ^43^) at the genus level did not identify significantly different genera between mice colonized with the isoDCA+ and isoDCA-consortia (**Figure S2C**). Assessment of alpha diversity revealed non-significant differences in isoDCA+ and isoDCA-consortia colonized mice by both the Shannon and Simpson indices; analysis of beta diversity by Bray-Curtis dissimilarity revealed a clear overlap between the two groups (**Figure S2D**). Taken together, this data suggests that the composition of the microbiome of mice colonized with the isoDCA+ consortium and isoDCA-consortium are comparable.

To confirm that the engrafted consortia produced bile acids *in vivo* as expected, metabolomic analysis of fecal bile acids was performed. Colonization of mice with the isoDCA+ consortium resulted in significantly elevated fecal levels of isoDCA and increased conversion of DCA to isoDCA on day 35, compared to mice colonized with the isoDCA-consortium (**Figure 2E**). There were no differences in fecal TCA, CA, or DCA (**Figure S2E**), indicating there were no differences in substrate availability between groups. Additionally, there were no differences in the major PBAs (**Figure S2F**), other PBAs (**Figure S2G**), or SBAs (**Figure S2H**) analyzed. Overall, the isogenic isoDCA+ and isoDCA-consortia modulated fecal isoDCA levels as expected, while maintaining comparable bile acid pools between the groups.

After confirming that the isoDCA+ and isoDCA-consortia colonize AVNM-treated mice and produce bile acids as expected, we characterized Treg populations in intestinal tissues since isoDCA was reported to induce large intestine LP RORγt^+^Foxp3^+^ pTregs^21^. Cells were isolated from both the ileal and colonic LP on day 35 and immunophenotyped by flow cytometry. We observed induction of RORγt^+^Foxp3^+^ pTregs, as proportions of both Foxp3^+^ Tregs and of CD4^+^ T cells and by total cell number, in the ileal LP of mice colonized with the isoDCA+ consortium compared to mice colonized with the isoDCA-consortium (**Figure 2F**). An increase in the number of Foxp3^+^ Tregs was also observed in the ileal LP (**Figure 2G**). In the colonic LP, colonization with the isoDCA+ consortium induced RORγt^+^Foxp3^+^ pTregs, both as a proportion of Foxp3^+^ Tregs and of CD4^+^ T cells (**Figure 2H**). A near-significant increase in the proportion of Foxp3^+^ Tregs of CD4^+^ T cells was also observed in the colonic LP of isoDCA+ consortium-colonized mice (**Figure 2I**). IsoDCA+ consortium colonized mice also had increased numbers of RORγt^+^ type 3 innate lymphoid cells (ILC3s) and a decreased proportion of RORγt^+^ Th17s of CD4^+^ T cells in the ileal LP, but not in the colonic LP (**Figures S2I and S2J**). This data shows that Clostridia-derived isoDCA can modulate immunosuppressive cell populations in both the small and large intestine and may function to promote intestinal homeostasis.

### IsoDCA-mediated Treg induction is dependent on TGR5

Various SBA derivatives have been reported to modulate intestinal immune responses by signaling through a variety of nuclear and cell surface receptors including FXR, TGR5, RORγt^+^, and VDR^21–23,44–47^. Despite their similar overall structure, different bile acids can uniquely signal through these various receptors, resulting in divergent effects on different cell types^18,48^. One report suggested that isoDCA-mediated Foxp3^+^ Treg induction arose from antagonism of FXR in dendritic cells, which resulted in an anti-inflammatory transcriptional profile *in vitro*^21^. The receptors required for the induction of isoDCA-mediated Tregs *in vivo* have not been investigated. As a specialized SBA receptor, TGR5 may be a more relevant receptor for isoDCA than FXR since it has been linked to a broad spectrum of anti-inflammatory effects on both myeloid and T cell populations for DCA and LCA derivatives^45^. To delineate the contributions of both TGR5 and FXR to the induction of RORγt^+^Foxp3^+^ pTregs we colonized both *Gpbar1*(TGR5)^-/-^ and *Nr1h4*(FXR)^-/-^ mice (and heterozygous littermates) with our engineered consortia using the same ABX-depletion timeline as before (**Figure 3A**).

**Figure 3.**
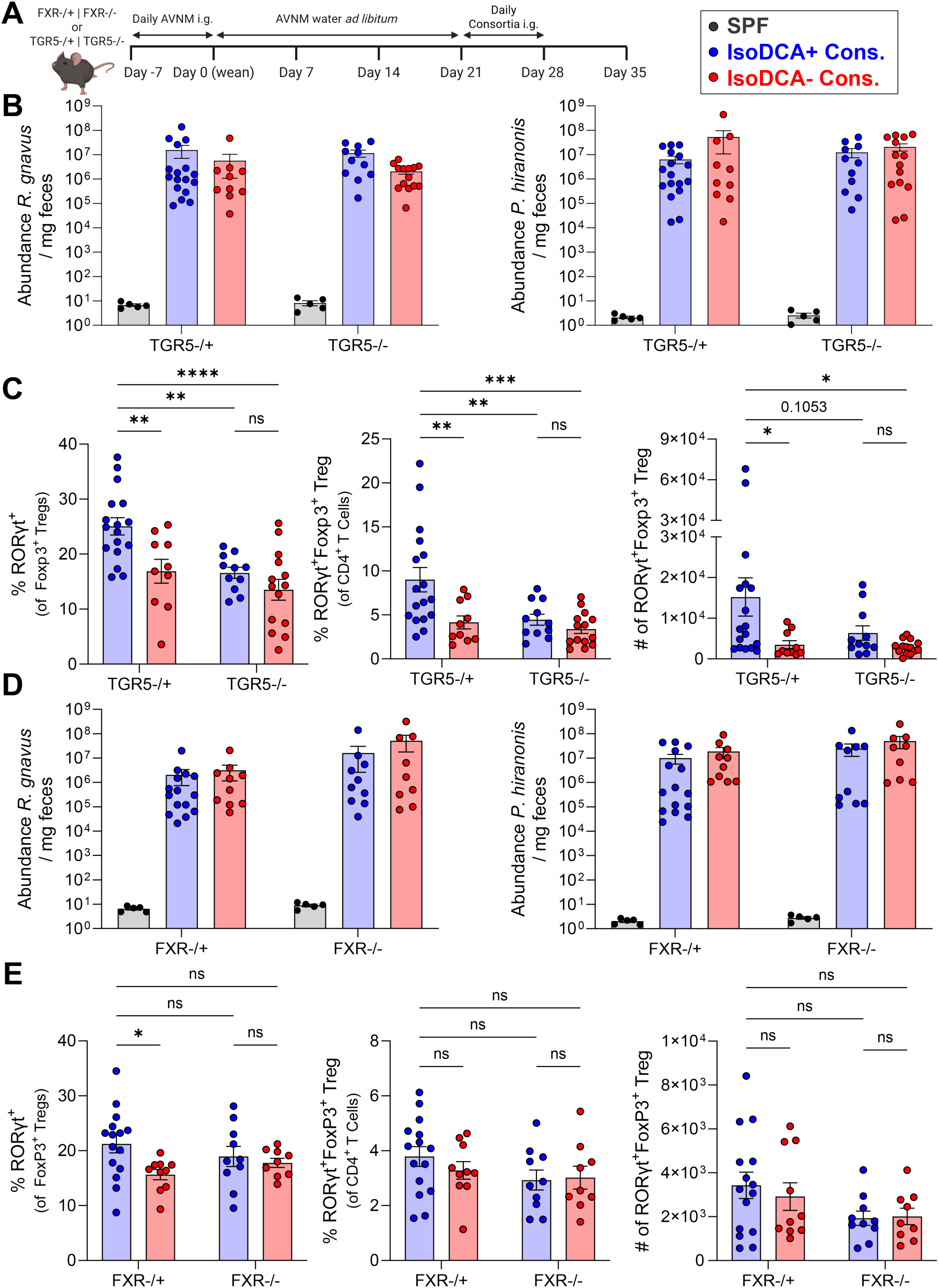
IsoDCA+ consortium-mediated colon lamina propria RORγt^+^FoxP3^+^ Treg induction is dependent on TGR5 and FXR. (A) Experimental timeline. (B) Abundance of *R. gnavus* (left) and *P. hiranonis* (right) normalized to fecal mass in TGR5^-/+^ and TGR5^-/-^littermates. (C) Proportion of RORγt^+^Foxp3^+^ Tregs of Foxp3^+^ Tregs (left) and of CD4^+^ T cells (center) and cell number of RORγt^+^Foxp3^+^ Tregs (right) in the colonic lamina propria (LP) of TGR5^-/+^ and TGR5^-/-^littermates on day 35. (D) Abundance of *R. gnavus* (left) and *P. hiranonis* (right) normalized to fecal mass in FXR^-/+^ and FXR^-/-^littermates. (E) Proportion of RORγt^+^FoxP3^+^ Tregs of FoxP3^+^ Tregs (left) and of CD4^+^ T cells (center) and cell number of RORγt^+^FoxP3^+^ Tregs (right) in the colonic LP of FXR^-/+^ and FXR^-/-^littermates on day 35. For (B, D), SPF = age-matched SPF mouse (*n* = 5). Data in (B, C) are pooled from four independent experiments. For (B, C), *n* = 17 isoDCA+ consortium in TGR5^-/+^, 10 isoDCA-consortium in TGR5^-/+^, 11 isoDCA+ consortium in TGR5^-/-^, 14 isoDCA-consortium in TGR5^-/-^. Data in (D, E) are pooled from four independent experiments. For (D, E), *n* = 15 isoDCA+ consortium in FXR^-/+^, 10 isoDCA-consortium in FXR^-/+^, 10 isoDCA+ consortium in FXR^-/-^, 9 isoDCA-consortium in FXR^-/-^. For (B-E), symbols represent individual mice and error bars represent mean ± SEM. For (C, E) statistics were analyzed by two-way ANOVA with Holm-Sidak’s post hoc test. * *P* < 0.05, ** *P* < 0.01, *** *P* < 0.001, **** *P* < 0.0001.

Engraftment was confirmed in *Gpbar1* (TGR5)^-/-^and *Gpbar1*(TGR5)^-/+^ mice on day 35 via qPCR for abundance of *P. hiranonis* and *R. gnavus* using species-specific primers (**Figure 3B**) and for fecal bacterial load using universal 16S primers (**Figures S3A and S3B**). Engraftment of the isoDCA+ consortium induced colonic LP RORγt^+^Foxp3^+^ pTregs in *Gpbar1*(TGR5)^-/+^ mice but not *Gpbar1*(TGR5)^-/-^mice (**Figure 3C**). RORγt^+^Foxp3^+^ pTregs were not induced in either *Gpbar1*(TGR5)^-/+^ or *Gpbar1*(TGR5)^-/-^mice engrafted with the isoDCA-consortium (**Figure 3C**), suggesting that TGR5 is necessary for isoDCA-mediated induction of colonic LP RORγt^+^Foxp3^+^ pTregs. No significant differences in colonic LP Foxp3^+^ Tregs were observed between *Gpbar1*(TGR5)^-/-^ and *Gpbar1*(TGR5)^-/+^ mice colonized with either the isoDCA+ or isoDCA-consortia (**Figure S3C**). *Gpbar1*(TGR5)^-/+^ mice colonized with the isoDCA+ consortium had increased proportions and numbers of RORγt^+^ ILC3s compared to isoDCA-consortium colonized mice (**Figure S3D**). Colonization of *Gpbar1*(TGR5)^-/+^ with the isoDCA+ consortium mice also induced RORγt^+^ Th17 cells compared to other groups (**Figure S3E**).

To examine a role for FXR we colonized *Nr1h4*(FXR)^-/-^ and *Nr1h4*(FXR)^-/+^ mice and confirmed engraftment on day 35 via qPCR for abundance of *P. hiranonis* and *R. gnavus* using species-specific primers (**Figure 3D**) and for fecal bacterial load using universal 16S primers (**Figure S3F**). In *Nr1h4*(FXR)^-/-^ mice, we did not observe differences in colonic LP RORγt^+^Foxp3^+^ pTreg populations between the groups engrafted with the isoDCA+ and isoDCA-consortia (**Figure 3E**). There was a significant increase in the proportion of colonic LP Foxp3^+^ Tregs which were RORγt^+^Foxp3^+^ in *Nr1h4*(FXR)^-/+^ mice colonized with the isoDCA+ consortium compared to those colonized isoDCA-consortium (**Figure 3E**). The increase in the proportion of colonic LP Foxp3^+^ Tregs which were RORγt^+^Foxp3^+^ in isoDCA+ consortium-colonized *Nr1h4*(FXR)^-/+^ compared to both consortia in the colonized *Nr1h4*(FXR)^-/-^ mice was not statistically significant (**Figure 3E**). Between all conditions, there were no differences in colonic LP Foxp3^+^ Tregs, RORγt^+^ Th17s, or RORγt^+^ ILC3s (**Figures S3G-S3I**) suggesting that the FXR-mediated effects of the isoDCA+ consortium were restricted to modulating the colonic LP pTreg compartment. It was previously reported that DC^ΔFXR^ mice (*Csf1r*^cre^*Nr1h4*^fl/fl^) had elevated large intestine LP RORγt^+^Foxp3^+^ pTregs compared to littermate DC^WT^ mice (*Csf1r*^WT^*Nr1h4*^fl/fl^)^21^. Thus, it is possible that in *Nr1h4*(FXR)^-/-^ mice, an aberrant BA pool and higher colonic LP pTregs at baseline are limiting any observed isoDCA-mediated RORγt^+^Foxp3^+^ induction. In the ileal LP, we did not observe isoDCA-mediated RORγt^+^Foxp3^+^ pTreg induction in *Gpbar1*(TGR5)^-/+^ or *Nr1h4*(FXR)^-/+^ mice (data not shown), possibly due to insufficient ileal isoDCA production. Overall, the evidence suggests that TGR5 is necessary for isoDCA-mediated induction of colonic LP RORγt^+^Foxp3^+^ pTregs, and that FXR may also contribute, albeit to a lesser extent.

### IsoDCA+ consortium attenuates intestinal inflammation

Given that the isoDCA+ consortium produced isoDCA and robustly induced immunosuppressive RORγt^+^Foxp3^+^ pTregs in the colonic LP it seemed possible that it could modulate T cell responses to promote intestinal homeostasis under inflammatory conditions. To interrogate this, we used the CD45RB^hi^ adoptive T cell transfer model of colitis where protection from disease can be mediated by co-transfer of Tregs that attenuate the differentiation, and subsequent pro-inflammatory activation, of transferred naïve CD4^+^ T cells^49,50^. SPF C57BL/6 *Rag1*^-/-^ mice were colonized with the isoDCA+ or isoDCA-consortium as before (**Figure 4A**). Two weeks after colonization, male and female *Rag1*^-/-^ mice received 4x10^5^ naïve CD4^+^ CD45RB^hi^ T cells with or without 5x10^4^ CD4^+^CD25^+^CD45RB^lo^ Tregs, a sub-optimal dose for complete protection, from female C57BL/6 donors (**Figure 4A**).

**Figure 4.**
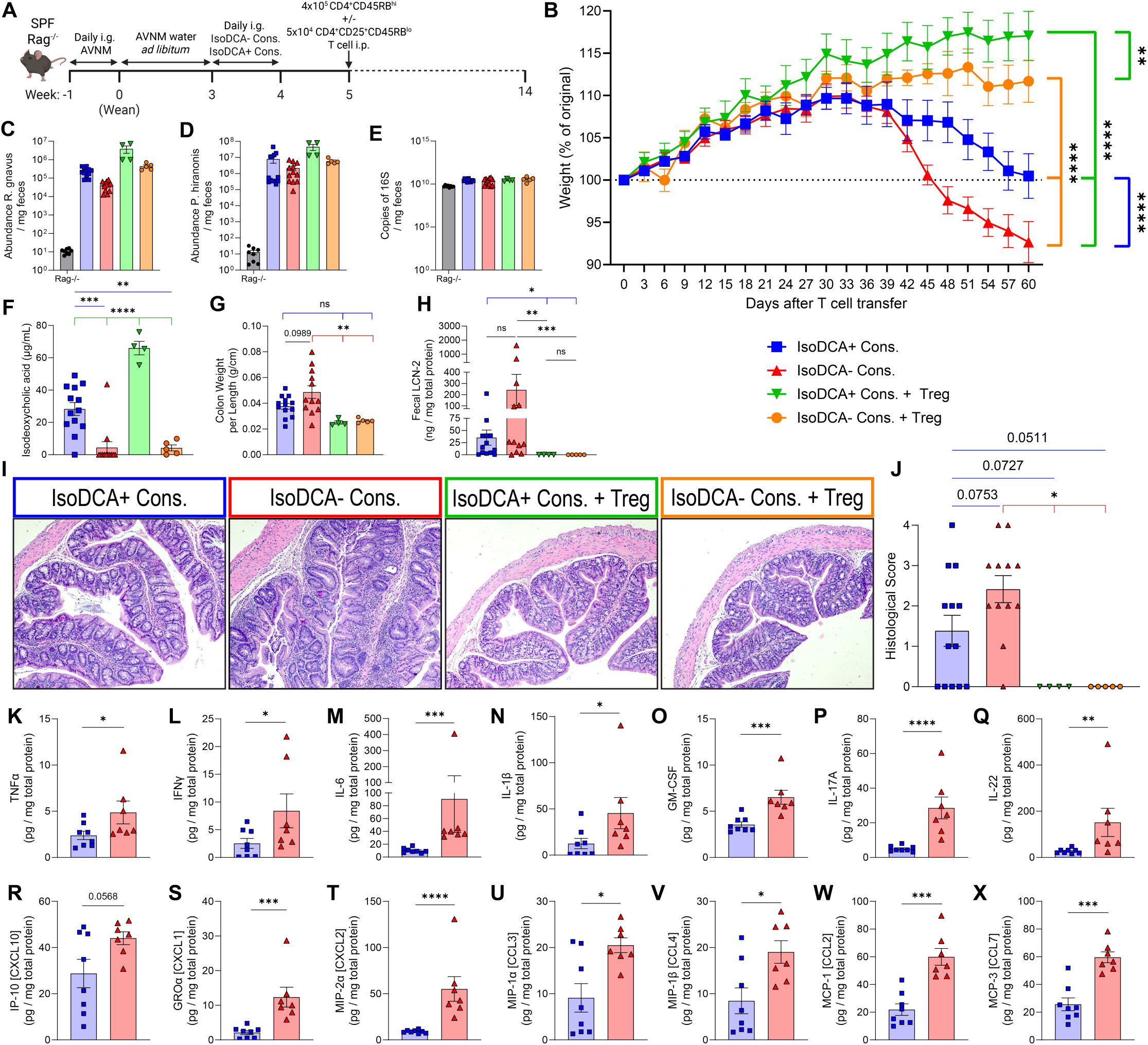
IsoDCA+ consortium protects against CD45RB^hi^ adoptive T cell transfer colitis and inhibits intestinal inflammation. (A) Experimental timeline. (B) Change in body weight as a percentage of starting body weight at time of T cell transfer. (C-D) Abundance of *R. gnavus* (C) and *P. hiranonis* (D) in feces collected day 60 post T cell transfer, normalized to fecal mass. (E) Fecal bacterial load measured on day 60 post T cell transfer, normalized to fecal mass. (F) Concentration of fecal isoDCA on day 60 post T cell transfer. (G) Weight to length ratio of colon tissue. (H) Lipocalin-2 (LCN-2) levels normalized to total protein concentration in feces collected on day 60 post T cell transfer. (I) Representative images of proximal colon tissues sections from each group stained with hematoxylin and eosin. Images were taken at 10X magnification. (J) Histological scoring of proximal colon tissue sections. (K-X) Concentration of TNFα (K), IFN-γ (L), IL-6 (M), IL-1β (N), GM-CSF (O), IL-17 (P), IL-22 (Q), IP-10 [CXCL10] (R), GROα [CXCL1] (S), MIP-2α [CXCL2] (T), MIP-1α [CCL3] (U), MIP-1β [CCL4] (V), MCP-1 [CCL2] (W), and MCP-3 [CCL7] (X) in lysed homogenates of middle colon sections normalized to total protein concentration. Data in (B-J) are pooled from two independent experiments. *n* = 13 isoDCA+ consortium, 12 isoDCA-consortium, 4 isoDCA+ consortium + Treg, and 5 isoDCA-consortium + Treg. Data in (K-X) are shown for one experimental repeat. *n* = 8 isoDCA+ consortium, 7 isoDCA-consortium. For (B), symbols represent mean ± SEM. Statistics were analyzed by area under the curve (AUC) one-way ANOVA with Tukey’s post hoc test. For (C-H, J-X), symbols represent individual mice and error bars represent mean ± SEM. For (F, G), statistics were analyzed by one-way ANOVA with Tukey’s post hoc test. For (H), statistics were analyzed by one-way ANOVA with Tukey’s post hoc test of log normalized values. For (K-X), statistics were analyzed by Student’s t-test of log normalized values. * *P* < 0.05, ** *P* < 0.01, *** *P* < 0.001, **** *P* < 0.0001.

To monitor disease progression, mice were weighed every 3 days for 60 days (**Figure 4B**). Weight loss is a non-invasive longitudinal readout of disease severity^50^. For groups that received only naïve CD4^+^ T cell transfer (standard transfer), colonization with the isoDCA+ consortium significantly protected against weight loss compared to colonization with the isoDCA-consortium (**Figure 4B**). Groups that received co-transfer of naïve CD4^+^ T cells and Tregs were largely protected from weight loss when compared to standard transfer as expected (**Figure 4B**). Interestingly, in co-transferred mice, isoDCA+ consortium colonized mice gained more weight than isoDCA-consortium-colonized mice suggesting that isoDCA may be enhancing the protective effects of co-transferred Tregs (**Figure 4B**). Broadly, across both the standard and co-transfer groups, the isoDCA+ consortium protected against weight loss when compared to isoDCA-consortium (**Figure 4B**). Consortium engraftment was confirmed at euthanasia on day 60 via qPCR for abundance of *P. hiranonis* and *R. gnavus* using species-specific primers and for fecal bacterial load using universal 16S primers (**Figures 4C-4E**).

16S rRNA sequencing on day 60 revealed that microbiota composition was similar between all experimental groups at the class level (**Figure S4A**). For the standard transfer groups, ALDEx2 (ref. ^43^) at the genus level identified that Gram-negative *Parasutterella* and unknown *Sutterellaceae* were significantly enriched in mice colonized with the isoDCA-consortium, and *Clostridioides* was significantly enriched in mice colonized with the isoDCA+ consortium (**Figure S4B**). It is unclear whether these genera are altered between groups due to direct effects of the isoDCA+ and isoDCA-consortia, or whether these genera are differentially regulated in the C57BL/6 *Rag1*^-/-^ mice due to immunodeficiency and/or the host inflammatory response and reflect the animals’ health status; these differentially abundant genera were not identified in consortium colonized C57BL/6 WT mice (**Figure S2C**). Alpha and beta diversity analysis showed similarities in microbiota composition between the standard transfer groups (**Figure S4C**). For mice that received Treg co-transfer, ALDEx2 identified more differentially abundant genera than for mice that only received naïve CD4^+^ T cell (standard) transfer (**Figure S4B**). IsoDCA+ consortium-colonized mice were enriched for *Sporofaciens* and unknown *Oscillospiraceae* and *Bacteroidales* genera (**Figure S4B**). Like standard transfer, Treg co-transfer isoDCA-consortium-colonized mice were enriched for Gram-negative taxa (*Paramuribaculum* and unknown *Muribaculaceae*), as well as *Adlercreutzia* and unknown *Prevotellaceae* (**Figure S4B**). Overall, the Shannon index is amplified in mice that received Treg co-transfer compared to mice that standard transfer (**Figure S4C**).

Fecal metabolomics revealed that, on day 60, mice colonized with the isoDCA+ consortium had elevated fecal isoDCA compared to mice that received the isoDCA-consortium and that of isoDCA+ consortium colonized mice, the Treg co-transfer group had elevated fecal isoDCA compared to the standard transfer group (**Figure 4F**). There were no significant differences in the upstream bile acids (**Figure S4D**), suggesting no differences in substrate availability. IsoDCA+ consortium colonized mice that received standard transfer had increased β-muricholic acid compared to Treg co-transfer, as well as isoDCA-consortium colonized standard transfer mice (**Figure S4E**). Additionally, isoDCA-consortium colonized mice that received Treg co-transfer had elevated fecal levels of lithocholic acid and several derivatives (**Figure S4E**). Notably, while no significant differences were observed in fecal levels of the immunomodulatory SCFAs butyrate and propionate; both Treg co-transfer groups had significantly elevated acetate compared to the standard transfer groups (**Figure S4F**).

In accordance with protection from weight loss, mice that received Treg co-transfer had lower colon weight to length ratio and fecal lipocalin-2 (LCN-2), a marker of neutrophil activity, compared to mice that received naïve CD4^+^ T cell (standard) transfer only (**Figures 4G and 4H**). Of the standard transfer colitic mice, those colonized with the isoDCA-consortium had the highest colon weight to length ratio and fecal LCN-2 (**Figures 4G and 4H**), indicating more severe colitis. Accordingly, histopathology revealed that for mice that received standard transfer, those colonized with the isoDCA-consortium had more severe lymphocytosis and epithelial architecture disruption compared to isoDCA+ consortium colonized mice (**Figures 4I and 4J**). Both Treg co-transfer groups were protected from lymphocytosis and epithelial architecture disruption (**Figures 4I and 4J**). Taken together, the data suggests that engraftment of the isoDCA secreting mini consortium mediates partial protection against colitis in the CD45RB^hi^ adoptive T cell transfer model.

Given that the isoDCA+ consortium induces immunosuppressive RORγt^+^Foxp3^+^ pTregs in the colonic LP (**Figure 2**), we speculated that the mechanism of protection would be isoDCA-mediated induction of colonic pTregs. However, for the standard transfer groups, immunophenotyping of colonic LP Treg populations isolated 60 days after T cell transfer revealed no significant differences in RORγt^+^Foxp3^+^ pTregs, Foxp3^+^ Tregs, and T-bet^+^Foxp3^+^ type 1 regulatory (Tr1) cells between isoDCA+ and isoDCA-consortium colonized mice (**Figures S4G-S4I**). It possible that over the course of the 60 days following T cell transfer, Treg populations equilibrated between groups and differences that were observed 14 days after colonization (**Figure 2I**) were no longer detectable. Campbell et al. reported isoDCA-mediated large intestine LP RORγt^+^Foxp3^+^ pTreg induction 10 days after colonization^21^. 4 weeks after colonization, differences in large intestine LP RORγt^+^Foxp3^+^ pTreg populations between isoDCA+ and isoDCA-groups (and the statistical significance) were diminished^21^. Taken together, this may suggest that initial exposure to isoDCA may promote a spike in RORγt^+^Foxp3^+^ pTreg induction which is lost over time. As expected, mice that received co-transfer of Tregs had increased populations of RORγt^+^Foxp3^+^ pTregs, Foxp3^+^ Tregs, and T-bet^+^Foxp3^+^ Tr1s compared to mice that received standard transfer (**Figures S4G-S4I**). A minor increase in the number of Foxp3^+^ Tregs and T-bet^+^Foxp3^+^ Tr1s was seen in the colonic LP of isoDCA-consortium-colonized mice with standard transfer compared to the co-transfer group (**Figures S4H and S4I**), which is likely due to broad expansion of T cells in the inflammatory context; broad expansion of Treg populations have previously been described in inflammatory conditions in this model^51,52^.

To further characterize the immunomodulatory effects of the isoDCA+ consortium under standard transfer conditions, we used a multiplex assay to examine cytokine production in homogenized colon lysates prepared at euthanasia. Compared to isoDCA+ consortium colonized mice, mice colonized with the isoDCA-consortium had elevated levels of pro-inflammatory cytokines TNFα, IFN-γ, IL-6, IL-1β, and GM-CSF (**Figures 4K-4O**), as well as elevated levels of type 3 cytokines IL-17A and IL-22 (**Figures 4P and 4Q**). By contrast mice colonized with the isoDCA-consortium had elevated levels of IP-10 [CXCL10] (an IFN-γ signaling marker and activated T cell recruitment chemokine), GROα [CXCL1] (neutrophil recruitment), MIP-2α [CXCL2] (neutrophil recruitment), MIP-1α [CCL3] and MIP-1β [CCL4] (related chemokines for macrophage, monocyte, and T cell recruitment), MCP-1 [CCL2] (monocyte, T cell, and dendritic cell recruitment), and MCP-3 [CCL7] (monocyte, eosinophil, basophil, and T cell recruitment) (**Figures 4R-4X**). No significant differences were observed for IL-2, IL-4, IL-5, IL-9, IL-10, IL12p70, IL-13, IL-18, IL-23, IL-27, RANTES [CCL5], or eotaxin [CCL11] (**Figure S4J**). All in all, this data suggests that engraftment with the isoDCA+ consortium protects against the CD45RB^hi^ adoptive transfer model of colitis by inhibiting pro-inflammatory cytokine signaling (mixed Th1/Th17 profile) and chemokine recruitment.

## Discussion

In this manuscript we describe a novel Clostridial mini consortium that we engineered to study the effects of isoDCA, an SBA previously shown to induce RORγt^+^Foxp3^+^ pTregs in the colonic LP^21^. This subset of pTregs has widely been recognized as central to promoting immunological tolerance to both dietary and commensal microbiota-derived antigens at mucosal surfaces^51–56^. RORγt^+^Foxp3^+^ pTregs have been reported to be protective in various murine models of NCCDs and have been clinically associated with tolerance to dietary antigens in humans^51–56^. To enable study of isoDCA produced *in situ* by native Clostridia we engineered *R. gnavus*, a commensal Clostridia that is widely found in humans^57^, to ablate its native isoDCA production. Although recent advancements have been made in Clostridial mutagenesis strategies, particularly for species in the genus *Clostridium*, non-model species remain largely recalcitrant to these strategies due to the highly polyphyletic nature of Clostridia. By disrupting *rumgna_00694*, which encodes the terminal enzyme in the biosynthetic pathway for isoDCA production, we created an isogenic strain, *R. gnavus* KO, with ablated isoDCA production (**Figures 1A-1D**). To better characterize our mutation, we complemented the system to ensure that targeted disruption of *rumgna_00694* was responsible for the observed ablation of isoDCA production (**Figures S1B and S1C**). Thus, we verified an isogenic approach to turn the conversion of DCA into isoDCA “on” or “off”.

To translate this approach to *in vivo* models, we needed to introduce BSH and *bai* operon functionalities to first convert host-derived TCA into DCA. Only a few Clostridial species have the *bai* operon necessary for 7α-dehydroxylation of PBAs^58^. *Clostridium scindens* and, to a lesser extent, *Clostridium hylemonae* are model species for 7α-dehydroxylation and remain the best characterized^19,58–61^. However, we instead selected *P. hiranonis*, a Clostridia known to uniquely possess both a BSH and the *bai* operon required to convert the host-derived conjugated PBA TCA into the SBA DCA ^33–35,62^, as a sole “primary degrader” of TCA to avoid adding in a third species for BSH activity and increasing the complexity of our system. Thus, we created the isoDCA+ consortium (*P. hiranonis* and *R. gnavus* WT) which can convert TCA to isoDCA *in vitro* and the isoDCA-consortium (*P. hiranonis* and *R. gnavus* KO) as an isogenic control with ablated isoDCA production (**Figure 1F**) to facilitate *in vivo* investigations into how isoDCA promotes intestinal homeostasis.

Because bile acid metabolism by the microbiota is a perpetual process in the lower GI tract, the engineered Clostridial consortia needed to recreate this constant metabolic flux to accurately model the physiological effects of isoDCA. To accomplish this on-going production, we successfully engrafted the Clostridial consortia in antibiotic-treated SPF mice (**Figures 2B-2D**). Mice colonized with the isoDCA+ consortium had elevated fecal isoDCA, and increased conversion of DCA to isoDCA, compared to isoDCA-consortium-colonized mice (**Figure 2E**). Importantly, this approach yielded isoDCA produced *in situ* by native Clostridia from physiologically relevant concentrations of host-derived TCA. In these colonized mice, we confirmed that isoDCA can induce both colonic LP RORγt^+^Foxp3^+^ pTregs (**Figure 2H**) and ileal LP RORγt^+^Foxp3^+^ pTregs (**Figure 2F**). Taken together, our data suggests that isoDCA, produced *in situ* by native Clostridia, can promote intestinal homeostasis through the induction of immunosuppressive RORγt^+^Foxp3^+^ pTregs in a biologically relevant context.

We then used the engineered Clostridial consortia to interrogate the *in vivo* mechanism of isoDCA-mediated RORγt^+^Foxp3^+^ pTreg induction. IsoDCA was previously suggested to promote *in vitro* Foxp3^+^ Treg induction through antagonism of FXR in DCs which yielded an anti-inflammatory phenotype^21^. We leveraged our novel Clostridial consortia to examine the roles of the bile acid receptors FXR and TGR5 *in vivo*. Using *Gpbar1*(TGR5)^-/-^ mice (and heterozygous littermates), we made the novel and unexpected finding that TGR5 is necessary for isoDCA mediated induction of colonic LP RORγt^+^Foxp3^+^ pTregs (**Figure 3C**). Colonization of *Nr1h4*(FXR)^-/-^ mice (and heterozygous littermates) with our mini consortium showed that FXR signaling may also play a role, although isoDCA-mediated FXR-dependent colonic LP RORγt^+^Foxp3^+^ pTreg induction was limited (**Figure 3E**).

Given the robust induction of colonic LP RORγt^+^Foxp3^+^ pTregs across several models, we then investigated whether isoDCA can protect against colitis in the CD45RB^hi^ adoptive T cell transfer model. In this model, transfer of naïve CD4^+^ T cells to immunodeficient *Rag1*^-/-^ mice results in a pro-inflammatory mixed Th1/Th17 effector T cell response specific to pathobionts, like *Helicobacter* spp., in the microbiota^49,63–65^. Notably, co-transfer of Tregs along with naïve CD4^+^ T cells protects against colitis, in a dose-dependent manner, by inhibiting the differentiation of effector T cells^49^. In these experiments, we found that the Clostridial consortia remained engrafted for over two months, and at the terminal endpoint, mice colonized with the isoDCA+ consortium maintained significantly elevated fecal isoDCA levels compared to isoDCA-consortium-colonized mice (**Figures 4C-4F**). IsoDCA+ consortium-colonized mice that received only naïve CD4^+^ T cells were protected from body weight loss and histopathological colonic inflammation compared to isoDCA-consortium-colonized mice that received only naïve CD4^+^ T cells (**Figures 4B, 4I, and 4J**), suggesting that the isoDCA+ consortium mediates protection from colitis. As expected, mice that received co-transfer of Tregs were protected from colitis compared to both groups of mice that only received transfer of naïve CD4^+^ T cells (**Figures 4B, 4I, and 4J**).

We did not detect increased numbers of colonic LP RORγt^+^Foxp3^+^ pTregs at euthanasia (9.5 weeks post T cell transfer and 11.5 weeks post colonization) in mice that received naïve CD4^+^ T cell (standard) transfer colonized with the isoDCA+ consortium compared to the isoDCA-consortium (**Figure S4G**); increased numbers of Foxp3^+^ Tregs were, however, detectable in both groups of mice that received Treg co-transfer (**Figure S4I**). In a subsequent experiment with naive CD4^+^ T cell (standard) transfer only, we characterized the immunosuppressive effects of the isoDCA+ consortium through multiplex analysis of homogenized colon lysates and found reduced proinflammatory chemokines and type 1 and type 3 cytokines compared to mice colonized with the isoDCA-consortium (**Figures 4K-4X**).

In summary, we created an engineered Clostridial mini consortium to interrogate the immunomodulatory effects of the secondary bile acid isoDCA. We leveraged these novel consortia to reveal new insights into the mechanism by which this prominent Clostridial metabolite promotes intestinal immune homeostasis. We demonstrated that isoDCA mediated induction of colonic LP RORγt^+^Foxp3^+^ pTregs, is dependent on the Takeda G-protein coupled receptor TGR5. Long term engraftment of the engineered isogenic mini consortia in mice with a replete SPF microbiome enabled examination of the biological function of a single microbial metabolite. We demonstrated that the isoDCA+ consortium (but not the isoDCA– consortium) protected against intestinal inflammation in the CD45RB^hi^ adoptive T cell transfer model by both modulating the composition of the microbiota and inhibiting pro-inflammatory responses.

## Methods

### Murine studies

Specific pathogen free (SPF) C57BL/6, *Rag1*^-/-^ (C57BL/6 background), *Gpbar1*(TGR5)^-/-^ (C57BL/6 background), and *Nr1h4*(FXR)^-/-^ (mixed background) mice were bred and housed at the University of Chicago Animal Resource Center. Mice were weaned at 21 days of age and were fed an irradiated diet (Teklad 2918, Inotiv). *Gpbar1*(TGR5)^-/-^ mice were provided by Dr. Kristina Schoonjans^44,46^ (École Polytechnique Fédérale de Lausanne, EPFL), and *Nr1h4*(FXR)^-/-^ mice were purchased from Jackson Labs. All experiments used both males and females and were littermate controlled. All mice used in these experiments were housed under a 12-hour light/dark cycle with pine-shaving bedding. Mice were euthanized by CO_2_ asphyxiation followed by secondary cervical dislocation or cardiac exsanguination. All experiments were approved by the Institutional Animal Care and Use Committee of the University of Chicago and performed in compliance with the University of Chicago Animal Care and Use Protocols.

To extract DNA for genotyping of *Gpbar1*(TGR5)^-/-^ and *Nr1h4*(FXR)^-/-^ mice (and heterozygous littermates), ear snips were digested overnight at 55 °C in a solution of 100mM Tris-HCl, 0.2% sodium dodecyl sulfate, 5mM ethylenediaminetetraacetic acid (EDTA), and 200 mM NaCl supplemented with 0.07 mg/mL Proteinase K. Debris was pelleted by centrifuging at 21300 x g for 15 minutes. DNA was precipitated by transferring supernatant to equal volume isopropyl alcohol. DNA was pelleted by centrifuging at 21300 x g for 7 minutes and resuspended in 200 μL UltraPure Distilled Water (Invitrogen). Genotyping was performed by polymerase chain reaction (PCR) using the primers in **Supplementary Table 1** and 2X HS Taq Mix (PCR Biosystems). Cycling conditions for *Gpbar1* (TGR5) genotyping are 95 °C for 3 minutes, followed by 38 cycles of 95 °C for 30 seconds, 59 °C for 30 seconds, and 72 °C for 30 seconds, and concluded with 72 °C for 10 minutes and an infinite hold at 4 °C. Cycling conditions for *Nr1h4* (FXR) genotyping are 94 °C for 5 minutes, followed by 35 cycles of 94 °C for 30 seconds, 60 °C for 30 seconds, and 72 °C for 30 seconds, and concluded with 72 °C for 7 minutes and an infinite hold at 4 °C. All genotyping PCR products were run on a 3% agarose gel at 110 V for 70 minutes.

### Metabolomic analysis of bile acids and short chain fatty acids

Metabolomic analyses of bile acids and short chain fatty acids in fecal samples were performed by the Duchossois Family Institute Host-Microbe Metabolomics Facility at the University of Chicago, as described previously^66^. Fecal samples were collected fresh and stored at -80 °C prior to submission. Fecal samples were weighed prior to submission.

### Clostridial engineering

To target the bile acid gene *rumgna_00694* in *R. gnavus* (VPI C7-9, originally acquired from ATCC (29149), group II introns were cloned into plasmid pJHA275 using previously described methods^32^. NCBI GenBank Accession AAYG00000000.2 was used for *R. gnavus*. Briefly, a re-targeted intron was designed using the ClosTron intron design tool ^29^ using the Perutka method ^67^ for the disruption of the 5’ coding DNA sequence of *rumgna_00694*. This re-targeted intron was subsequently cloned via Gibson assembly ^68^ into pJHA275 resulting in plasmid pJHA342. Transfer of the group II intron plasmid to *R. gnavus* was performed using previously described methods^32^. Briefly, genetic material was transferred from donor *Escherichia coli* S17 λpir□ to *R. gnavus* using conjugative mating strategies adapted from ref. ^69^. Prior to mating, plates containing BHI agar (BD Difco) supplemented with 0.05% L-cysteine-HCl (Sigma) were reduced in the anaerobic chamber for at least 4 hours. Saturated overnight cultures in selective media of *E. coli* containing pJHA342 were diluted 1:100 in fresh selective media and grown for approximately 4 hours to reach OD_600_ of 0.4-0.6. Concurrently, saturated overnight cultures of *R. gnavus* were diluted 1:100 and grown for approximately 4 hours to reach OD_600_ of 0.4-0.6. Once reached, 1 mL of donor *E. coli* culture was centrifuged for 5 minutes at 5000 x g. The supernatant was discarded, and the donor pellet was transferred to an anaerobic chamber (Coy Laboratory Products) where it was washed twice with anaerobic PBS containing 0.05% cysteine-HCl (with centrifugation at 5000 x g for 5 minutes). After washing, 250 μL of *R. gnavus* recipient culture was used to resuspend the *E. coli* donor pellet, and the entire suspension was added to reduced BHI plates supplemented with 0.05% cysteine-HCl in a single spot and rested at room temperature for at least 30 minutes to dry. Mating plates were then incubated anaerobically at 37 °C for 18-24 hours. Following incubation, mating spots were scraped using sterile 1 μL inoculation loops into 500 μL of anaerobic PBS and suspended by aspiration. After a homogeneous suspension was achieved, 250 μL of the suspended mating was plated on reduced selective BHI plates supplemented with 0.05% cysteine-HCl and containing 5 μg/mL thiamphenicol (Sigma) and 60 μg/mL kanamycin (Sigma). Plates were incubated at 37 °C for 24-72 hours until colonies appeared. Individual colonies were selected using a sterile inoculation loop and streaked onto fresh reduced selective BHI plates to isolate single colonies. Isolation plates were incubated anaerobically at 37 °C for 24-72 hours and isolated colonies were grown in reduced BHI media supplemented with 0.05% cysteine-HCl anaerobically at 37 °C for 24-72 hours. Colony PCR with GOTaq2 (NEB) was used to confirm the successful transfer of genetic material.

### PCR and qPCR of bacteria

Bacterial DNA was isolated from cultured bacterial samples or mouse fecal pellets using the DNeasy PowerSoil Pro (Qiagen) kit according to manufacturer’s protocol. For cultured samples, 1mL of culture was pelleted by centrifuging at 1500 x g for 5 minutes, aspirated, and resuspended in solution CD1 from the kit. For fecal samples, the weight of 1-2 pellets per mouse was recorded and transferred to solution CD1. Final DNA was eluted in a volume of 100 μL.

The presence of bacteria in either cultured bacterial or fecal samples was confirmed and measured via PCR using the primers in **Supplementary Table 2** using Dream Taq Polymerase (Thermo Scientific). DNA was extracted as described above. For detection of *rumgna_00694* (to differentiate *R. gnavus* WT and KO), cycling conditions are 95 °C for 2 minutes, followed by 35 cycles of 95 °C for 30 seconds, 61 °C for 30 seconds, and 72 °C for 2.5 minutes, and concluded with 72 °C for 5 minutes and an infinite hold at 4 °C. For quicker analysis of cultured samples, colony PCR was performed instead of DNA extraction. For this, 1mL of culture was pelleted by centrifuging at 1500 x g for 5 minutes. After aspiration, a small amount of pellet was picked up on a pipette tip and transferred to PCR reaction mix. PCR was run as described above, but with initial denaturation step extended to 10 minutes at 95 °C.

The abundance of bacteria in fecal samples was quantified via qPCR on a QuantStudio 3 Real-Time PCR System (ThermoFisher) using the primers in **Supplementary Table 2** using the PowerUp SYBR Green Master Mix Kit (AppliedBiosystems). DNA was extracted as described above. For measuring abundances of *R. gnavus* and *P. hiranonis*, assays were run using the comparative Ct (ΔΔ Ct) experiment type. For *R. gnavus* the PCR stage conditions are 35 cycles of 95 °C for 15 seconds, 56 °C for 30 seconds, and 72 °C for 20 seconds. For *P. hiranonis* the PCR stage conditions are 35 cycles of 95 °C for 30 seconds, 51 °C for 30 seconds, and 72 °C for 50 seconds. For both species, the Hold Stage is 50 °C for 2 minutes then 95 °C for 10 minutes, data is collected during the elongation step of the cycles, and the Melt Curve Stage is the default for SYBR reagents. For bacterial load quantification, the standard curve experiment type was used, and the cycling conditions are 40 cycles of 95 °C for 30 seconds, 52 °C for 30 seconds, and 72 °C for 60 seconds. Standard curves were generated from single copy 16S *Porphyromonas* sp. standard at 10^7.5^ copies/μL diluted to 10^7^, 10^6.5^, 10^6^, etc. The Hold Stage is 50 °C for 2 minutes then 95 °C for 2 minutes, data is collected during the elongation step of the cycles, and the Melt Curve Stage is the default for SYBR reagents. Bacterial DNA samples were diluted 1:1000 in UltraPure DNase/RNase-Free Water (Invitrogen) for bacterial load qPCRs.

To quantify abundance based on fecal mass per sample, the following equation was used: 2 ⁄. The values were very small, so they were multiplied by ∼10^13^ to normalize control groups close to 1. Limit of detection was determined by Ct of DNA extraction control, no template (water) control, or maximum thermocycles of the qPCR.

### Bacterial culture

*R. gnavus* WT, *R. gnavus* KO (*R. gnavus* Ω*rumgna_00694*), and *P. hiranonis* were cultured at 37 °C in an incubator inside a flexible vinyl anaerobic chamber with concentrations of H_2_ = 2.5-3.0% and O_2_ < 50 ppm. A scrubber for H_2_S was continuously used in the anaerobic chamber. For liquid culturing, bacteria were grown in supplemented brain heart infusion (BHI-S) medium consisting of 37 g/L of brain heart infusion powder (BD Difco), 5 g/L yeast extract (BD Bacto), and 1% (v/v) of 10% (w/v) L-cysteine (Sigma). L-cysteine was 0.22 μm sterile filtered and added to cooled, autoclave-sterilized BHI-S medium. For *in vitro* bile acid metabolism experiments, yeast casitone fatty acids broth with carbohydrates (YCFAC) medium (Anaerobe Systems) was used to avoid sterol compounds found in BHI-S medium. For agar plates, 15 g/L agar (Sigma) was included to BHI-S medium prior to autoclaving. All liquid media and plates were allowed to reduce in the anaerobic chamber overnight prior to use.

To make glycerol stocks, individual strains were streaked on BHI-S agar plates and allowed to grow for 48-96 hours or until individual colonies could be picked and cultured in 7 mL liquid BHI-S medium at 37 °C overnight or until turbid. Primary cultures were then used to inoculate secondary cultures in BHI-S medium that were grown at 37 °C for 12-16 hours to OD_600_ ≈ 0.800. Dilution plating was used to determine that OD_600_ = 0.800 correlated to ∼2 x 10^8^ CFU/mL for each stain. For glycerol stocks, 10 mL of *P. hiranonis* culture (OD_600_ = 0.800) was mixed with 10 mL of *R. gnavus* WT or KO culture (OD_600_ = 0.800) and pelleted by centrifuging at 2500 x g for 10 minutes. Resulting pellet was aspirated and resuspended in 1mL sterile BHI-S medium supplemented with 15% (v/v) sterile glycerol and stored at -80 °C.

For *in vitro* bile acid metabolism experiments, bacteria were cultured as described above by streaking on BHI-S plates and inoculating primary cultures in ∼7 mL YCFAC medium. For experiments investigating conversion of DCA, 3oxo-DCA, or TCA, secondary YCFAC cultures were spiked with 8 μL of 100 mM bile acids prior to reduction in the anaerobic chamber. Secondary cultures spiked with bile acids were inoculated with 50 μL of *P. hiranonis*, *R. gnavus* WT, or *R. gnavus* KO turbid primary cultures and grown at 37 °C overnight prior to liquid-liquid extraction. For some experiments investigating conversion of TCA, secondary cultures were spiked as described above and cultured with 50 μL of *P. hiranonis* turbid primary culture and grown at 37 °C overnight. *P. hiranonis* secondary cultures were spun at 2500 x g for 5 minutes then 0.22 μm sterile filtered. Sterile spent media from *P. hiranonis* secondary cultures were then inoculated with 50 μL of either *R. gnavus* WT or KO turbid primary cultures and grown at 37 °C overnight prior to liquid-liquid extraction.

### Liquid-liquid extraction and thin layer chromatography of bile acids

Liquid-liquid extraction of bile acids from bacterial culture was adapted from ref. ^21^. First, 250 μL 12 N HCl and 2 g NaCl were first added to 5 mL of culture and mixed by shaking. Then 4 mL ethyl acetate (EtOAc) was added and mixed well with regular venting. Layers were allowed to settle, the resulting organic (top) layer was collected, and the process was repeated with an additional 4 mL EtOAc. For the resulting ∼8 mL EtOAc extractant, 2 g anhydrous MgSO_4_ was added to remove residual water and filtered out using a 40 μm filter. The anhydrous EtOAc extractant was dried under vacuum centrifugation (Thermo Savant DNA120) in a fume hood. Dried pellets were resuspended with 100 μL methanol. Controls of sterile YCFAC medium neat and spiked with bile acids were extracted.

For thin layer chromatography (TLC), a beaker sealed with aluminum foil was used as the TLC chamber, with a solvent of 70:20:2 benzene:1,4-dioxane:glacial acetic acid^21^. A Kimwipe saturated with solvent was placed along the back of the beaker to equilibrate vapor pressure within the TLC chamber. Using glass-backed TLC Silica gel 60 F_254_ plates (Sigma), 2 μL of extracted samples and 0.5 μL of 10 mM control bile acid solutions in methanol were spotted onto plates. The solvent was allowed to run ∼90% of the plate length prior to plate removal and air drying for 5 minutes. After drying, plates were stained with 10% (w/v) copper (II) sulfate and 8% (v/v) phosphoric acid in deionized water then baked at 135 °C for 15 minutes prior to imaging.

### Bacterial complementation

For inducible gene expression, cloning was performed in an inducible expression plasmid (pJHA272) as previously described^32^. Briefly, a plasmid containing *rumgna_00694* under the control of an anhydrotetracycline-inducible promoter was assembled into pJHA272 using Gibson assembly to replace the existing NanoLuc coding region, creating pJHA826. This construct was transferred to *R. gnavus* KO (*R. gnavus* Ω*rumgna_00694*), to restore the 3β-HSDH function of *R. gnavus*.

To assess *in vitro* bile acid production by complementation, cultures of *R. gnavus* WT and *R. gnavus* KO, *R. gnavus* KO + *rumgna_00694* (*R. gnavus* KO comp.) and *R. gnavus* KO + empty vector control (*R. gnavus* KO empty) were cultured using BHI-S plates and YCFAC medium spiked with 60 μg/mL kanamycin (Sigma) and 5 μg/mL thiamphenicol (Sigma) to prevent growth of *E. coli* or *R. gnavus* KO that shed their plasmid, respectively. Secondary cultures of YCFAC medium were also spiked with bile acids, as described in *Liquid-liquid extraction and thin layer chromatography of bile acids*, with the addition of 1 μg/mL anhydrotetracycline (ThermoFisher) to induce gene expression. Bile acids were extracted and TLC was performed as before.

### Colonization of mice with Clostridial consortia

SPF C57BL/6, *Rag1*^-/-^, *Gpbar1*(TGR5)^-/-^ (and heterozygous littermates), and *Nr1h4* (FXR)^-/-^ (and heterozygous littermates) mice were bred and housed at the University of Chicago. At 14 days old, pups began antibiotic treatment by daily intragastric (i.g.) gavage of 100 μL PBS supplemented with 25 mg/mL ampicillin sodium salt (Sigma), 1.25 mg/mL vancomycin hydrochloride (Sigma), 25 mg/mL neomycin trisulfate salt hydrate (Sigma), and 5 mg/mL metronidazole (Sigma). On day 21 of age, pups were weaned onto ampicillin (1 g/L), vancomycin (0.5 g/L), neomycin (1 g/L), and metronidazole (0.2 g/L) supplemented acidified drinking water in water bottles with no lixit for 3 weeks. After 3 weeks, antibiotic water was removed and the lixit and an acidified water bottle were replaced. Immediately following the removal of antibiotic, mice began colonization by daily i.g. gavage of 200μL bacterial glycerol stocks of the isoDCA+ consortium (*P. hiranonis* and *R. gnavus* WT) or the isoDCA-consortium (*P. hiranonis* and *R. gnavus* KO) for 7 days. For the first 3 days of bacterial glycerol stock gavage, mice were housed in new cages after gavage to prevent antibiotic carry-over by coprophagy.

### 16S rRNA sequencing and analysis

16S rRNA sequencing and analysis was performed by the Duchossois Family Institute Microbiome Metabolomics Facility (DFI MMF) at the University of Chicago. Fecal samples were collected fresh and stored at -80 °C prior to submission. Weighed fecal samples or extracted DNA were submitted for analysis. For submitted fecal samples, DFI MMF extracted DNA using the QIAamp PowerFecal Pro DNA kit (Qiagen). For all samples, the V4-V5 region of 16S rRNA genes were PCR amplified using barcoded dual-index primers. Illumina compatible libraries were generated using the Qiagen QIASeq 1-step amplicon kit and sequencing was performed on the DFI MMF Illumina MiSeq platform using a 2×250 Paired End reads, generating 5,000-10,000 reads per sample. Raw V4-V5 16S rRNA gene sequence data was demultiplexed and processed through the dada2 pipeline into Amplicon Sequence Variants (ASVs). ASVs were identified with the Bayesian RDP classifier up to the genus level and were BLASTed against RefSeq for species-level identification with corresponding alignment statistics.

All analyses of 16S sequencing data were performed in R. Taxonomic classification and sequence data were integrated using the phyloseq and Biostrings packages. ASVs were processed from .fasta files and mapped to classification metadata. Taxonomic bar plots were visualized at the class level using ggplot2. Taxa falling outside the top 15 most abundant features were categorized as “Other” and shaded in grey to emphasize dominant community shifts. Plots were faceted by experimental group and displayed with relative abundance percentages.

Differential abundance analysis was conducted using the ALDEx2 (Analysis of different abundance taking sample and scale variation into account) package^43^. Data were filtered to remove low-prevalence taxa (present in <2 samples) prior to analysis. The ALDEx2 pipeline employed a Dirichlet-multinomial model to generate 128 Monte Carlo instances, which were then centered log-ratio (clr) transformed. Significantly differentially abundant taxa were identified using a Wilcoxon rank-sum test threshold of 0.05 and a standardized effect size cutoff of 1.0.

Alpha diversity was evaluated using the Shannon and Simpson indices. For two-group comparisons, statistical significance was determined using the non-parametric Wilcoxon rank-sum test. For multi-group plots, a global Kruskal-Wallis rank-sum test was first employed to identify overall differences in diversity. Post-hoc pairwise Wilcoxon tests were subsequently performed to resolve significant differences between specific group pairs, with an alpha of 0.05.

Beta diversity was visualized using Principal Coordinates Analysis (PCoA) based on Bray-Curtis dissimilarity. Prior to ordination, counts were transformed into relative abundance using total sum scaling. Ninety-five percent confidence ellipses were calculated for each group to visualize community dispersion.

### Lamina propria (LP) lymphocyte isolation for flow cytometry

Intestinal epithelial cells were first removed from ileal (defined as the final 10 cm of small intestine) and colonic tissue. Following tissue harvest and the removal of mesenteric fat, tissues were rinsed with PBS, cut open longitudinally, and stored in PBS on ice. Peyer’s patches were not removed. Tissues were transferred to 4 mL of ice-cold PBS supplemented with 30mM EDTA and 1.5mM dithiothreitol and incubated on ice for 20 minutes, followed by a subsequent incubation at 37 °C in PBS + 30mM EDTA for 10 minutes. IECs were mechanically dissociated from tissue by physical shaking for ∼30 seconds. The remaining tissue was transferred to 4mL ice cold PBS prior to digestion.

To isolate LP lymphocytes, tissues were first mechanically dissociated by chopping with surgical scissors in 4mL RPMI-1640 (Cytiva) supplemented with 4% fetal bovine serum (FBS) (Cytiva), 50 μg/mL LiberaseTM (Roche), and 40 μg/mL DNase I (Sigma), followed by enzymatic digestion at 37 °C in an orbital shaker for 40 minutes, with vigorous shaking by hand every 10 minutes. Cell suspensions were passed through 70 μm sterile nylon filters prewet with PBS supplemented with 4% FBS, tubes and filters were washed with 4% FBS/PBS, and cell suspension was pelleted by centrifuging at 800 x g for 5 minutes. Supernatant was aspirated and resuspended in 4% FBS/RPMI-1640. 4 mL of Percoll (Cytiva) diluted to 80% in PBS with a final concentration of 1X was mixed with the cell suspension (to create a 40% Percoll cell suspension) and overlayed on 5 mL of 80% Percoll to create a 40% / 80% Percoll interface. Samples were centrifuged at 1200 x g for 35 minutes with acceleration set to 1 and brake set to 0. Lymphocytes were collected from the interface into 10 mL 4% FBS/RPMI. Cells were pelleted by centrifuging at 800 x g for 5 minutes. Following a wash with 1mL 4% FBS/RPMI-1640, supernatant was carefully aspirated and cells were resuspended in 250 μL 4% FBS/RPMI for cell counting. To count, 10 μL of cells were mixed with 10 μL of trypan blue (Invitrogen) and counted via Countess II (Invitrogen).

### Flow cytometry

For flow cytometric analysis of LP lymphocytes, single cell suspensions were plated into round bottom 96 well plates and washed (pelleted by centrifuging at 800 x g for 3 minutes, aspirated, and resuspended in 200 μL) with PBS. Cells were stained for viability via Fixable Live/Dead Aqua or Fixable Live/Dead Blue (ThermoFisher) by resuspending in 25 μL of stain diluted 1:500 in 1X PBS on ice for 15 minutes in the dark. Cells were washed first with PBS then with 1% bovine serum albumin (BSA) in PBS. Fc receptor signaling was blocked by incubating cells on ice in the dark for 10 minutes by resuspending cells in 25 μL of anti-mouse CD16/CD32 (InVivoMAb) at a final concentration of 0.1 mg/mL in 1% BSA/PBS. Cells were then stained for extracellular staining of CD3ε, CD4, CD25, Lineage markers, CD45, CD90, or CD45RB using the antibodies in **Key Resources Table** by adding 25 μL to wells and incubating for 30 minutes at room temperature in the dark. Extracellular stain was prepared by first adding antibodies to 5X Brilliant Stain Buffer Plus (BD Biosciences) then adding 1% BSA/PBS to dilute to final concentration. Cells were washed twice with 1% BSA/PBS.

For intracellular staining, cells were fixed using the FOXP3/Transcription Factor Staining Buffer Set (eBioscience). Cells were incubated with 100 μL of the fixing solution (prepared according to instructions) overnight at 4°C. Cells were washed with 1% BSA/PBS followed by permeabilization buffer (prepared according to instructions). For intracellular staining, cells were first resuspended in 25 μL permeabilization buffer, then incubated with 25 μL intracellular stain for Foxp3, RORγt, or T-bet using antibodies listed in the **Key Resources Table** for 30 minutes at room temperature in the dark. Intracellular stain for transcription factors was prepared similarly to the extracellular stain with antibodies first being added to Brilliant Stain Buffer Plus but diluted in permeabilization buffer to final concentration. Following staining, cells were washed twice with permeabilization buffer, once with 1% BSA/PBS and resuspended in 100 μL 1% BSA/PBS for analysis on the Aurora spectral flow cytometer (CytekBio). Cell populations were analyzed using FlowJo (BD). All flow cytometric analyses included florescence minus one (FMO) stained cells and single stained cells as controls.

### CD45RB^hi^ transfer model of murine colitis

SPF *Rag1*^-/-^ mice were colonized with the isoDCA+ or isoDCA-consortia as before. After ANVM drinking water was removed and colonization gavages began, mice were housed in cages containing soiled bedding from segmented filamentous bacteria-positive and *Helicobacter*-positive *Rag1*^-/-^ mice. Two weeks after colonization began (35 days post-weaning) mice were injected intraperitoneally (i.p.) with 4 x 10^5^ CD45RB^hi^CD4^+^CD25^-^cells with or without 5 x 10^4^ CD45RB^lo^CD4^+^CD25^+^ cells in 100 μL PBS to induce colitis^50^. Donor cells were isolated from spleens of 6–8-week-old female donor SPF C57BL/6 mice via enzymatic digestion, as described for intestinal tissues, for 30 minutes at 37°C prior to filtering through 70 μm sterile nylon filters. CD4^+^ cells were enriched from pooled splenocytes using EasySep™ Mouse CD4+ T Cell Isolation Kit (Stemcell Technologies) according to manufacturer’s protocol. Enriched CD4^+^ cells were stained for flow cytometry using the protocol described above, adapted for 5mL flow cytometry tubes by washing with 500 μL and staining with a final volume of 100 μL. Enriched CD4^+^ cells were only stained extracellularly with Fixable Live/Dead Blue and CD4, CD25, and CD45RB antibodies. CD45RB^hi^CD4^+^CD25^-^cells (defined as the top 40% of CD45RB expressing cells ^70^ and CD45RB^lo^CD4^+^CD25^+^ cells were sorted using a Bigfoot (ThermoFisher) cell sorter into tubes with final concentrations of 5% (v/v) FBS, 1 mM EDTA, 10 mM HEPES, and 100 U/mL penicillin and streptomycin. Isolated cells were washed twice with PBS, counted, and diluted to a final concentration of 4 x 10^6^ CD45RB^hi^CD4^+^CD25^-^cells/mL with or without 5 x 10^5^ CD45RB^lo^CD4^+^CD25^+^ cells/mL in PBS.

Prior to i.p. transfer of T cells, all recipient mice were weighed. Mice were monitored daily and body weight was recorded every 3 days to measure disease progression. Body weight measurements were taken around the same time to minimize circadian effects. The experiments were terminated when mice lost ∼20% of starting body weight (∼60 days after T cell transfer). Colon was harvested and the tissue length and weight (contents removed, flushed with PBS) were recorded. Proximal, middle, and distal colon sections were collected for histological analysis.

For one experiment, LP cells were isolated from the remaining colon tissue and entire ileum tissue for flow cytometric analysis. For the second experiment, remaining colon tissue was cut into ∼1cm sections (noting the order from proximal to distal colon) and snap frozen in tubes by submerging in isopropanol cooled by dry ice for multiplex cytokine/chemokine analysis.

### Histology of mouse colon

Tissue sections (∼0.5 cm) of the proximal, middle, and distal colon were harvested and fixed in 10% buffered formalin phosphate at 4 °C overnight. Sections were processed, paraffin embedded, cut into 5 μm cross-sections, mounted on slides, and stained with hematoxylin and eosin (H&E) by the University of Chicago Human Tissue Resource Center. H&E-stained proximal colon samples were blindly evaluated and scored as in **Supplementary Table 3** by Christopher Weber M.D. Ph.D., a GI pathologist at UChicago Medicine. Samples were imaged with a Zeiss Axiocam 506 at 10X magnification.

### Measurement of fecal lipocalin-2 (LCN-2) by ELISA

Fecal samples were collected from mice and analyzed fresh or stored at -80°C until analysis. Samples were weighed and homogenized in 0.5 mL of PBS supplemented with 0.1% (v/v) Tween 20. To homogenize, samples were vortexed for 10 minutes then centrifuged at 12000 x g at 4 °C for 10 minutes to pellet debris. Supernatant was collected, aliquoted, and used fresh for subsequent analysis. Total protein was quantified by Pierce BCA Protein Assay Kit (Thermo Scientific) according to manufacturer’s protocol and using PBS + 0.1% Tween 20 as the diluent. Fecal LCN-2 levels were quantified using Mouse Lipocalin-2/NGAL DuoSet ELISA (R&D Systems) according to manufacturer’s protocol. BCA protein assays and ELISAs were analyzed with a SpectraMax M3 reader (Molecular Devices). Fecal LCN-2 levels were normalized to total fecal protein concentration.

### Measurement of intestinal cytokines by multiplex immunoassay

Colon sections were harvested and snap frozen. For multiplex immunoassay, two ∼1 cm sections of middle colon tissue were weighed and homogenized together in 400 μL ProcartaPlex Cell Lysis Buffer (ThermoFisher) in 2mL Safe-Lock tubes (Eppendorf) by bead beating using a 5 mm stainless steel ball bearing (Qiagen) at 25 Hz for 3 minutes on a Retsch MM 400 mixer. Cell debris was pelleted by centrifuging at 16000 x g for 10 minutes at 4 °C. Supernatants were collected and aliquoted. Total protein concentration was quantified by Pierce BCA Protein Assay Kit (Thermo Scientific) according to manufacturer’s protocol. Cytokines and chemokines were quantified using the ProcartaPlex Mo Cytokine/Chemokine Panel 1 26plex (ThermoFisher). Assay was performed according to manufacturer’s instructions with sample binding performed overnight at 4 °C. Samples were run on a Luminex 200. Cytokine and chemokines levels were normalized to total protein concentration for analysis. Samples were analyzed fresh, and remaining lysates were aliquoted and frozen at -80°C.

## Supporting information

Graphical Abstract

## Acknowledgments

We thank Dr. Kristina Schoonjans (École Polytechnique Fédérale de Lausanne) for providing *Gbpar1*(TGR5)^-/-^mice. We would also like to thank the Duchossois Family Institute (DFI) and the University of Chicago Cytometry and Antibody Technology Facility for their assistance with DNA sequencing and flow cytometry, respectively. BioRender was used for diagrams. This work was supported by NIH grants R21AI186046 (C.R.N.) and R35GM147478 (M.M.) and an NSF Graduate Research Fellowship (J.H.A.).

## Author Contributions

EI and CRN designed the study. JA designed the plasmids and generated *R. gnavus* KO with input from MM. EI and CRN wrote the manuscript. C.R.W. performed and scored the histopathology. All authors read and commented on the manuscript.

## Declaration of Interest

The authors declare no competing interests.

## Data Availability

Data supporting the findings of this study are available within the article, its supplementary information, and the accompanying Source Data files. 16S rRNA sequencing data is deposited at NCBI under accession number PRJNA1503769. Other data are provided in the accompanying Source Data. Bacterial strains are available upon request upon contacting the corresponding author.

## Notes

### Competing Interest Statement

The authors have declared no competing interest.

