## Supplementary Information for "An engineered Clostridial mini consortium modulates intestinal inflammation"

**Table S1. Murine primers.** List of primers for murine genotyping PCR.

| Primer Name | Primer Sequence (5’ -> 3’) | Reference |
|---|---|---|
| FXR R | GCA TGC TCT GTT CAT AAA CGC CAT | Jackson Labs Protocol: 23042 |
| FXR WT F | TCT CTT TAA GTG ATG ACG GGA ATC T | Jackson Labs Protocol: 23042 |
| FXR KO F | GCT CTA AGG AGA GTC ACT TGT GCA | Jackson Labs Protocol: 23042 |
| TGR5_rec3 | AGA TGC TTT GGG TAG GTT GC | Pols et al *Cell Metabolism,* 2011 |
| TGR5_ko3_common | GGT GGG TGA GTG GAG TCT TC | Pols et al *Cell Metabolism*, 2011 |
| TGR5_WT3 | ACG AGT GCT TCG AGG AG | Pols et al *Cell Metabolism,* 2011 |

**Table S2. Bacterial primer sequences.** List of primers for analysis of bacterial taxa.

| Primer Name | Assay | Primer Sequence (5’ -> 3’) | Reference |
|---|---|---|---|
| H35 (*rumgna_00694* F) | PCR | TGT TGG CTT GGC ATT ACC GTG G | This study |
| H36 (*rumgna_00694* R) | PCR | TGC ACA ATT TTC CAG ATG CCG GC | This study |
| Phira F (*P. hiranonis* F) | qPCR | AGT AAG CTC CTG ATA CTG TCT | Ohashi Y et al, *Biosci Microbiota Food Health. 2019* |
| Phira R (*P. hiranonis* R) | qPCR | GGG AAA GAG GAG ATT AGT CC | Ohashi et al, *Biosci Microbiota Food Health. 2019* |
| Rg16S F (*R. gnavus* F) | qPCR | TGG CGG CGT GCT TAA CA | Joossens M et al, *Gut*, 2019 |
| Rg16S R (*R. gnavus* R) | qPCR | TCC GAA GAA ATC CGT CAA GGT | Joossens M et al, *Gut*, 2019 |
| 8F (Universal 16S F) | qPCR | AGA GTT TGA TCC TGG CTC AG | Turner S et al, J Eukaryot Microbiol. 1999 |
| 338R (Universal 16S R) | qPCR | TGC TGC CTC CCG TAG GAG T | Amann et al, Microbiol Rev. 1995 |

**Table S3. Histopathological scoring.** Scoring criteria for histopathological analysis.

| Score | Description |
|---|---|
| 0 | Without diagnostic abnormality |
| 1 | Patchy clusters of increased lymphocytes near bases of crypts |
| 2 | Diffuse increased lymphocytes in lamina propria and epithelium |
| 3 | Diffuse increased lymphocytes in lamina propria and epithelium, and with patchy crypt drop out, lamina propria fibrosis, or some granulomatous inflammation |
| 4 | Marked lymphocytosis with architectural distortion and some crypt abscesses |

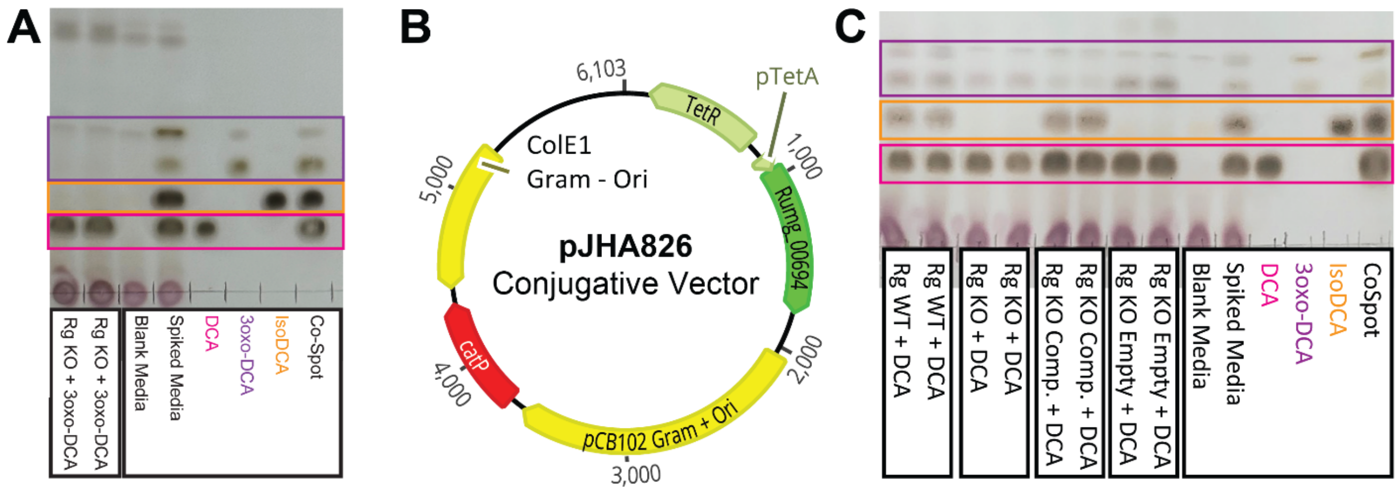


**Figure S1.**
(A) Representative thin layer chromatography (TLC) of culture supernatants of *R. gnavus Ωrumgna_00694* (Rg KO) supplemented with 3oxo-DCA.
(B) Plasmid used to induce *rumgna_00694* expression in Rg KO for complementation.
(C) Representative thin layer chromatography of culture supernatants of *R. gnavus* WT (Rg WT), Rg KO, complemented Rg KO (Rg KO Comp.) and empty vector Rg KO (Rg KO Empty) supplemented with DCA.

For (A, C), repeat lanes indicate biological replicates.

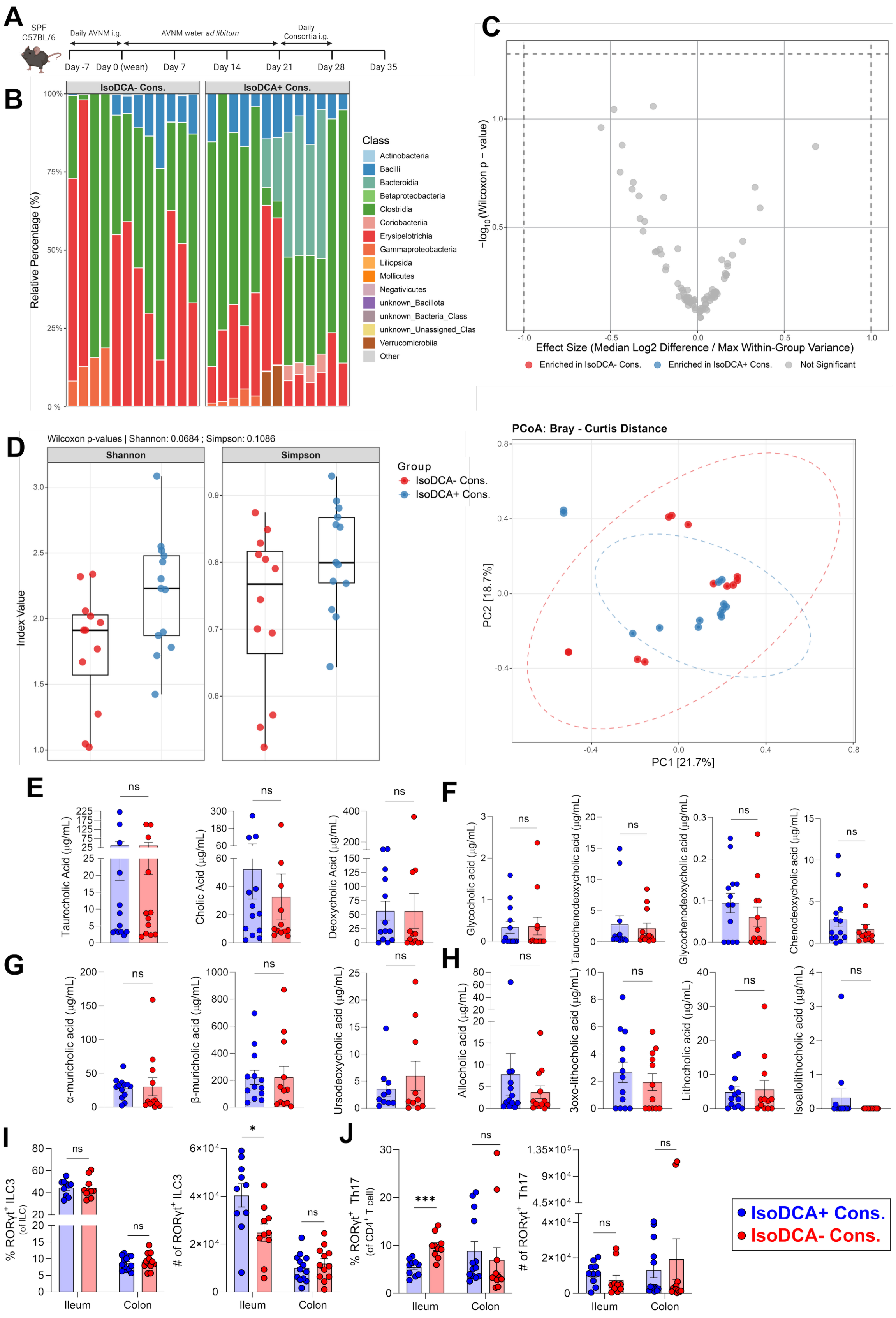


**Figure S2.**

(A) Experimental timeline.

(B) Composition of the microbiota on day 35 in mice colonized with the isoDCA- consortium (left) or the isoDCA+ consortium (right) from 16S rRNA sequencing, shown by bar plots at the Class level. Classes outside the top 15 are pooled in “Other”.

(C) Differentially abundant genera between mice colonized with the isoDCA+ or isoDCA- consortium by ALDEx2 (Analysis of different abundance taking sample and scale variation into account).
(D) Fecal microbiome Shannon and Simpson alpha (left, box plots) and beta (right, PCoA) diversity of mice colonized with the isoDCA+ or isoDCA- consortium. Dashed ellipses indicate 95% confidence.
(E) Fecal concentrations of taurocholic acid, cholic acid, and deoxycholic acid on day 35.
(F) Fecal concentrations of other primary bile acids: glycocholic acid, taurochenodeoxycholic acid, glycochenodeoxycholic acid, and chenodeoxycholic acid on day 35.
(G) Fecal concentrations of other bile acids: α-muricholic acid, β-muricholic acid, and ursodeoxycholic acid on day 35.
(H) Fecal concentrations of other secondary bile acids: allocholic acid, lithocholic acid, 3oxo-lithocholic acid, and isoallolithocholic acid on day 35.
(I) Proportion of RORγt$^+$ type 3 innate lymphoid cells (ILC) of ILCs (left) and number of RORγt$^+$ ILC3s (right) in the ileal and colonic lamina propria (LP) on day 35.
(J) Proportion of RORγt$^+$ T helper 17 (Th17) cells of CD4$^+$ T cells (left) and number of RORγt$^+$ Th17 cells (right) in the ileal and colonic LP on day 35.

Data in (B-L) are pooled from two independent experiments. For (B-D), $n$ = 13 isoDCA+ consortium, 12 isoDCA- consortium. For (E-H), $n$ = 13 isoDCA+ consortium, 12 isoDCA- consortium. For (I,J), ileal LP $n$ = 10 isoDCA+ consortium, 10 isoDCA- consortium and colonic LP $n$ = 13 isoDCA+ consortium, 12 isoDCA- consortium. For (B, D), bars and symbols represent individual mice, respectively. For (E-L), symbols represent individual mice and error bars represent mean ± SEM. Statistics were analyzed for each metabolite (E-H) and tissue (I, J) by Student's t-test. * $P < 0.05$, *** $P < 0.001$.

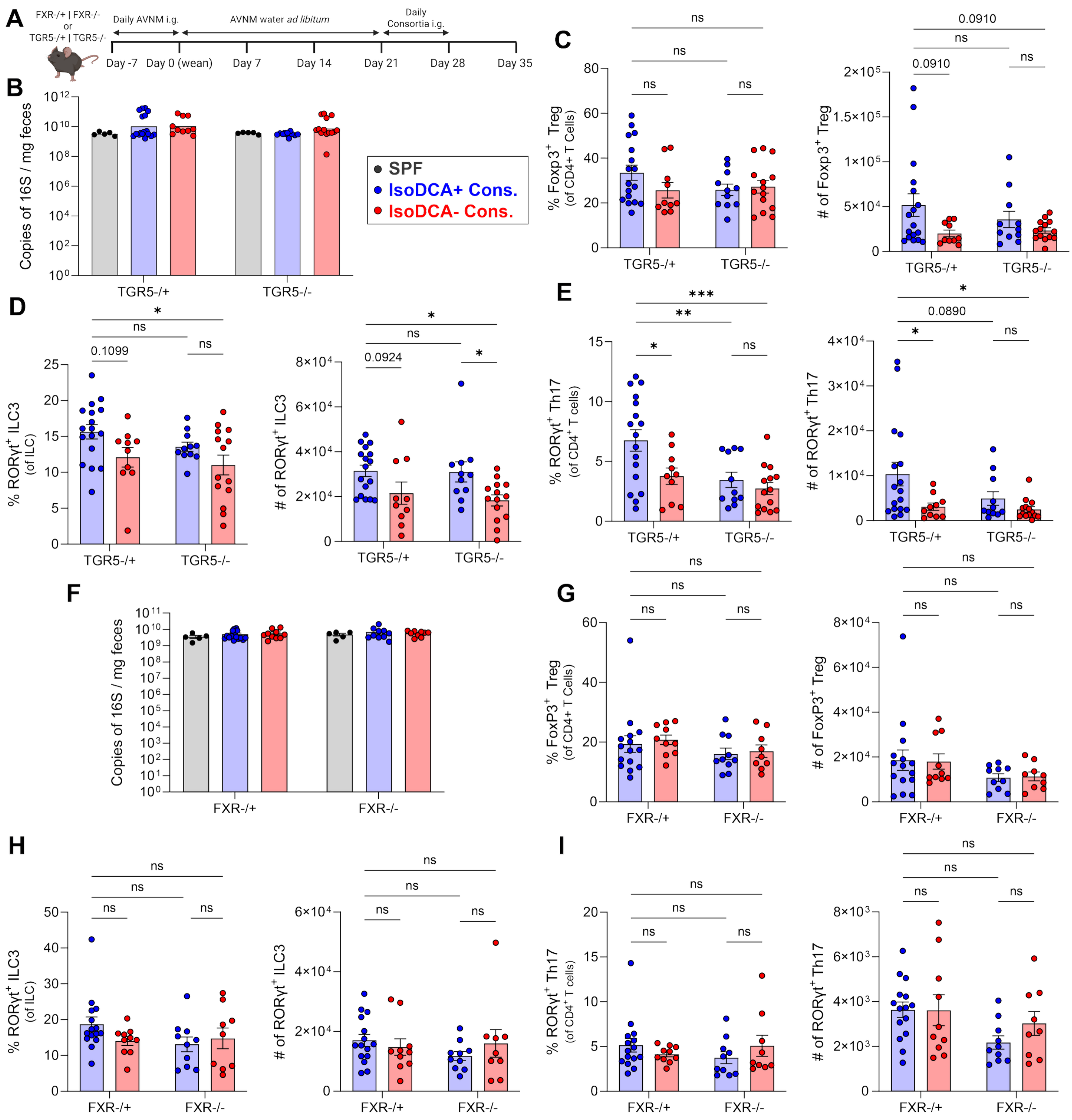


**Figure S3.**

(A) Experimental timeline.

(B) Fecal bacterial load normalized to fecal mass in TGR5$^{-/+}$ and TGR5$^{-/-}$ littermates.

(C) Proportion of Foxp3$^{+}$ Tregs of CD4$^{+}$ T cells (left) and number Foxp3$^{+}$ Tregs (right) in the colonic lamina propria (LP) of TGR5$^{-/+}$ and TGR5$^{-/-}$ littermates on day 35.

(D) Proportion of RORγt$^{+}$ type 3 innate lymphoid cells (ILC3) of ILC (left) and number of RORγt$^{+}$ ILC3 (right) in the colonic LP of TGR5$^{-/+}$ and TGR5$^{-/-}$ littermates on day 35.

(E) Proportion of RORγt$^{+}$ Th17 of CD4$^{+}$ T cells (left) and number of RORγt$^{+}$ Th17 (right) in the colonic LP of TGR5$^{-/+}$ and TGR5$^{-/-}$ littermates on day 35.

**Figure S3. continued.**
(F) Fecal bacterial load normalized to fecal mass in $FXR^{-/+}$ and $FXR^{-/-}$ littermates.
(G) Proportion of $Foxp3^{+}$ Tregs of $CD4^{+}$ T cells (left) and number of $Foxp3^{+}$ Tregs (right) in the colonic LP of $FXR^{-/+}$ and $FXR^{-/-}$ littermates on day 35.
(H) Proportion of $ROR\gamma t^{+}$ ILC3 of ILC (left) and number of $ROR\gamma t^{+}$ ILC3 (right) in the colonic LP of $FXR^{-/+}$ and $FXR^{-/-}$ littermates on day 35.
(I) Proportion of $ROR\gamma t^{+}$ Th17 of $CD4^{+}$ T cells (left) and number of $ROR\gamma t^{+}$ Th17 (right) in the colonic LP of $FXR^{-/+}$ and $FXR^{-/-}$ littermates on day 35.

For (B, F), SPF = level of age-matched SPF mouse (*n* = 5). Data in (B-E) are pooled from four independent experiments. For (B-E), *n* = 17 isoDCA+ consortium in $TGR5^{-/+}$, 10 isoDCA- consortium in $TGR5^{-/+}$, 11 isoDCA+ consortium in $TGR5^{-/-}$, 14 isoDCA- consortium in $TGR5^{-/-}$ mice. Data in (F-I) are pooled from four independent experiments. For (F-I), *n* = 15 isoDCA+ consortium in $FXR^{-/+}$, 10 isoDCA- consortium in $FXR^{-/+}$, 10 isoDCA+ consortium in $FXR^{-/-}$, 9 isoDCA- consortium in $FXR^{-/-}$ mice. For (B-I), symbols represent individual mice and error bars represent mean ± SEM. For (C-E, G-I) statistics were analyzed by two-way ANOVA with Holm-Sidak's post hoc test.

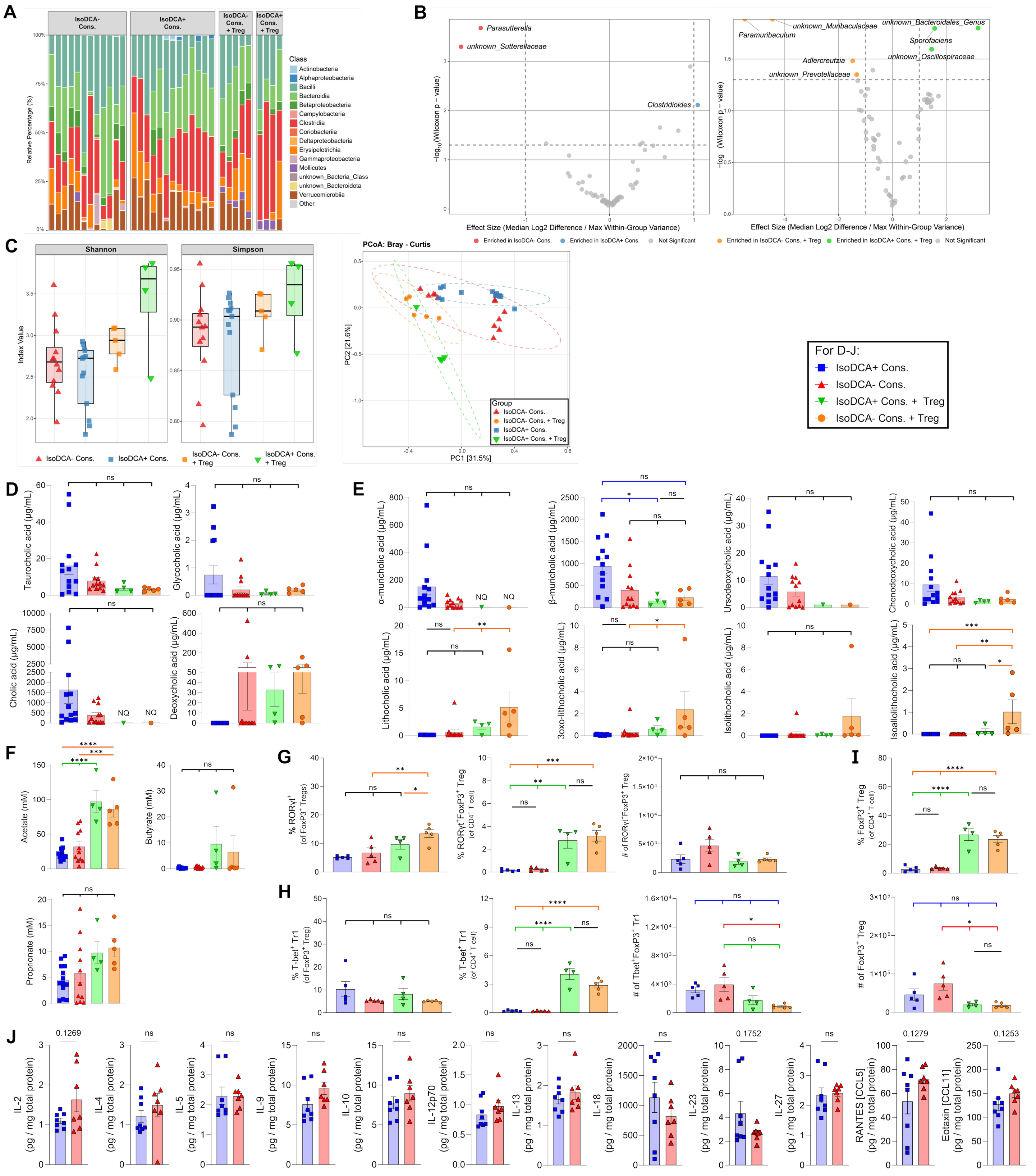


**Figure S4.**
(A) Microbiota composition on day 60 after CD4$^+$CD45RB$^{hi}$ T cell transfer in mice colonized with the isoDCA- consortium with or without Tregs (left) or the isoDCA+ consortium with or without Tregs (right) from 16S rRNA sequencing, shown by bar plots at the Class level. Classes outside the top 15 are pooled in “Other”.

(B) Differentially abundant genera between mice colonized with the isoDCA+ or isoDCA- consortium without Tregs (left) and with Tregs (right) by ALDEx2 (Analysis of different abundance taking sample and scale variation into account).
(C) Fecal microbiome Shannon and Simpson alpha diversity of mice colonized with the isoDCA+ or isoDCA- consortium without Tregs (left) and with Tregs (center) and Bray Curtis beta diversity (right, PCoA). Dashed ellipses indicate 95% confidence.
(D) Fecal concentrations of taurocholic acid, glycocholic acid, cholic acid, and deoxycholic acid on day 60 post T cell transfer. NQ = not quantified.
(E) Fecal concentrations of other bile acids: α-muricholic acid, β-muricholic acid, ursodeoxycholic acid, chenodeoxycholic acid, lithocholic acid, 3oxo-lithocholic acid, isolithocholic acid, and isoallolithocholic acid on day 60 post T cell transfer. NQ = not quantified.
(F) Fecal concentrations of short chain fatty acids: acetate, butyrate, and propionate on day 60 post T cell transfer.
(G) Proportion of RORγt$^+$Foxp3$^+$ Tregs of Foxp3$^+$ Tregs (left) and of CD4$^+$ T cells (center) and number of RORγt$^+$Foxp3$^+$ Tregs (right) in the colonic LP 60 days after T cell transfer.
(H) Proportion of T-bet1$^+$Foxp3$^+$ type 1 regulatory cells (Tr1) of Foxp3$^+$ Tregs (left) and of CD4$^+$ T cells (center) and number of T-bet1$^+$Foxp3$^+$ Tr1s (right) in the colonic LP 60 days after T cell transfer.
(I) Proportion of Foxp3$^+$ Tregs of CD4$^+$ T cells (top) and number of Foxp3$^+$ Tregs (bottom) in the colon LP 60 days after T cell transfer.
(J) Concentration of IL-2, IL-4, IL-5, IL-9, IL-10, IL-12p70, IL-13, IL-18, IL-23, IL-27, RANTES [CCL5], and Eotaxin [CCL11] in lysed homogenates of middle colon sections normalized to total protein concentration.

Data in (A-F) are pooled from two independent experiments. For (A-E), *n* = 13 isoDCA+ consortium, 12 isoDCA- consortium, 4 isoDCA+ consortium plus Treg, and 5 isoDCA- consortium plus Treg. For (F), *n* = 13 isoDCA+ consortium, 11 isoDCA- consortium, 5 isoDCA+ consortium plus Treg, and 5 isoDCA- consortium plus Treg. For (A, C) bars and points represent individual mice, respectively. Data in (G-I) are shown for one experiment. *n* = 5 isoDCA+ consortium, 5 isoDCA- consortium, 4 isoDCA+ consortium plus Treg, and 5 isoDCA- consortium plus Treg. Data in (J) are shown for one experiment. *n* = 8 isoDCA+ consortium, 7 isoDCA- consortium. For (D-I), symbols represent individual mice and error bars represent mean ± SEM. Statistics were analyzed by one-way ANOVA with Tukey's post hoc test. For (J), statistics were analyzed by Student's t-test of log normalized values. * $P < 0.05$, ** $P < 0.01$, *** $P < 0.001$, **** $P < 0.0001$.
