## Supplementary figures and images for "An engineered Clostridial mini consortium modulates intestinal inflammation"

### Graphical Abstract

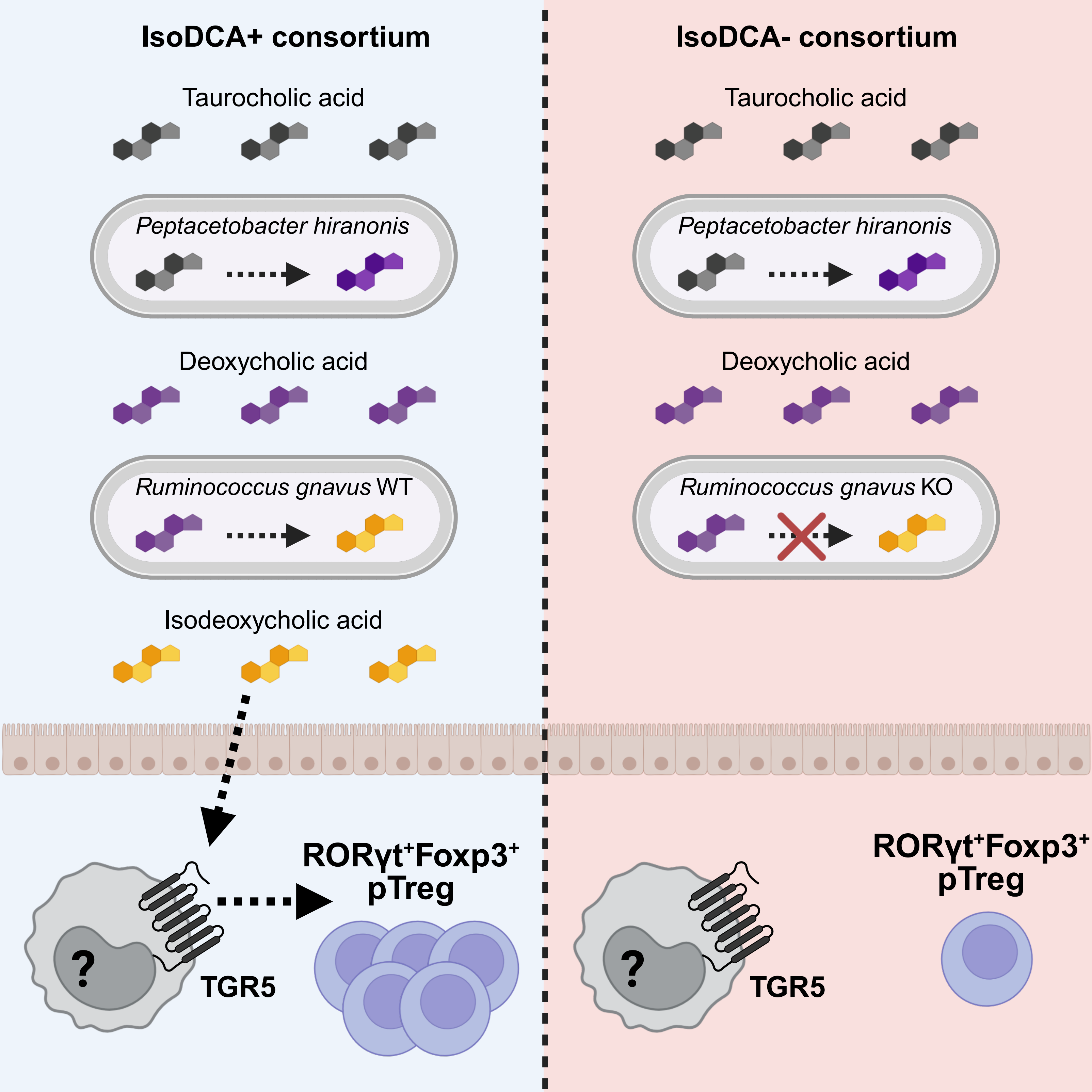
